# Synaptic architecture of a memory engram in the mouse hippocampus

**DOI:** 10.1101/2024.04.23.590812

**Authors:** Marco Uytiepo, Yongchuan Zhu, Eric Bushong, Filip Polli, Katherine Chou, Elise Zhao, Christine Kim, Danielle Luu, Lyanne Chang, Tom Quach, Matthias Haberl, Luca Patapoutian, Elizabeth Beutter, Weiheng Zhang, Brian Dong, Elle McCue, Mark Ellisman, Anton Maximov

## Abstract

Memory engrams are formed through experience-dependent remodeling of neural circuits, but their detailed architectures have remained unresolved. Using 3D electron microscopy, we performed nanoscale reconstructions of the hippocampal CA3-CA1 pathway following chemogenetic labeling of cellular ensembles with a remote history of correlated excitation during associative learning. Projection neurons involved in memory acquisition expanded their connectomes via multi-synaptic boutons without altering the numbers and spatial arrangements of individual axonal terminals and dendritic spines. This expansion was driven by presynaptic activity elicited by specific negative valence stimuli, regardless of the co-activation state of postsynaptic partners. The rewiring of initial ensembles representing an engram coincided with local, input-specific changes in the shapes and organelle composition of glutamatergic synapses, reflecting their weights and potential for further modifications. Our findings challenge the view that the connectivity among neuronal substrates of memory traces is governed by Hebbian mechanisms, and offer a structural basis for representational drifts.

## INTRODUCTION

In both natural and artificial systems, information is typically encoded within the structure of a carrier. It is widely accepted that the brain acquires long-term memories through the restructuring of neural circuits. This conceptual framework draws on mounting empirical evidence, indicating that sensory stimuli elicit morphological changes in neurons and individual synapses across brain regions essential for associative, spatial, and motor learning ^1–4^. However, current insights into the nature and origins of experience-dependent events that configure the physical substrates of memory traces are largely based on light imaging ^1–4^. Despite ongoing technical advancements, light imaging methods are unsuitable for detailed dissection of remarkably complex synaptic networks due to their limited resolution and capacity to simultaneously monitor distinct structural features. Hence, the architectures of circuits involved in memory storage remain poorly understood. This gap in knowledge hinders the elucidation of phenomena underpinning alternative models of neural computations for cognitive tasks.

The classic Hebbian theory, encapsulated by the “fire together, wire together” axiom, postulates that learning arises from a selective increase in synaptic weights between connected neurons consequent to their correlated excitation ^5^. Hebbian mechanisms are thought to manifest through the strengthening of existing synapses or through *de novo* synaptogenesis, leading to enhanced connectivity ^5–7^. The hypothesis that memories are stored within stable subsets of neurons gained support from pioneering studies that harnessed the immediate early gene *Fos* to identify and trace cellular ensembles recruited for learning ^8^. Subsequently, targeted optogenetic reactivation of these ensembles has been shown to be sufficient for memory retrieval without external cues, whereas their inhibition disrupted normal retrieval ^2,9–11^. The use of *Fos*-inducible fluorescent markers for optical imaging has also reinforced the idea that neurons coactivated during learning tend to form new synapses among themselves ^12,13^. Yet, the Hebbian model cannot easily be reconciled with recent discoveries demonstrating that cellular representations of sensory stimuli drift over time ^14–19^. This phenomenon implies that engram ensembles, even within the same brain area, might incorporate neurons that were not initially engaged. It is conceivable that representational drifts occur due to non-trivial re-patterning of synaptic connectivity, deviating from traditional Hebbian principles, but the underlying mechanisms have remained elusive.

In the present work, we sought to address these problems by using Serial Block Face Scanning Electron Microscopy (SBEM), a technique for three-dimensional (3D) nanoscale reconstruction of biological tissues ^20^. This technique permits the accurate alignment of sequentially obtained high resolution EM image planes to cover large fields of view without compression artifacts. Contemporary 3D-EM methods have been instrumental for deciphering the basic logic of neuronal wiring in *C.elegans*, *Drosophila* and different parts of the mammalian central nervous system, including the cerebral cortex, hippocampus, thalamus, spinal cord, and retina ^21–29^. Since saturated reconstructions of large 3D-EM stacks pose substantial challenges, most comprehensive connectomic analyses to date have presented data from a single specimen ^21,22,25,26,28^. Therefore, these studies do not explain how the intricate organization of the brain is influenced by the external world, internal state, or other factors such as age, sex, and genetic background. To attain our goal, we combined SBEM with behavior, chemogenetic tagging of physiologically relevant neurons, and artificial intelligence (AI) algorithms for image segmentation. This approach allowed us to explore the effects of distinct combinations of sensory cues at levels ranging from local connectomes to synaptic sub-compartments. Moreover, we were able to test the current models through quantitative measurements of various parameters in independent biological samples.

## RESULTS

### Permanent labeling of hippocampal engrams with APEX2-mGFP

We searched for structural correlates of long-term information storage in the hippocampus of mice subjected to contextual fear conditioning (CFC), a neurobehavioral paradigm for acquiring associative memory of an aversive environment where neutral visual and olfactory cues are paired with a brief series of unpleasant foot shocks ^2,8^. Given the need for sampling 3D images from multiple animals with a nanometer resolution, we focused on the Stratum Radiatum (sr) of area CA1, where dendrites of pyramidal glutamatergic neurons (PNs) are robustly innervated by Schaffer Collateral (SchC) axons of PNs residing in the upstream area CA3 (Figure 1A). The importance of the CA3-CA1 pathway for associative learning and memory has been well established ^30^. Considering that cellular ensembles recruited for each learning epoch are sparse ^2,8^, we devised a workflow for irreversible labeling of transiently activated populations with a genetically encoded engineered peroxidase, APEX2-mGFP (Figures 1B and S1). This plasma membrane-anchored, EM-compatible marker was introduced with the Cre recombinase-inducible adeno-associated virus (AAVDJ-DIO:APEX2-mGFP) into the hippocampus of previously characterized knock-in mice that express destabilized Cre (DD-Cre) under the control of the endogenous *Fos* promoter (*Fos^DD-Cre^*) ^31,32^. *Fos* is an immediate early gene whose transcription is restricted to narrow windows following neuronal excitation ^8,33^. The newly synthesized DD-Cre fusion proteins rapidly degrade through the proteasomal pathway, but their decay can be blocked by peripheral delivery of the blood-brain barrier (BBB)-permeable antibiotic, Trimethoprim (TMP) ^32,34^. Because free TMP crosses the BBB within minutes and also has fast kinetics of clearance from the brain ^32^, the *Fos^DD-Cre^* allele permits temporally restricted recombination of *LoxP*-flanked DNA sequences in neurons activated by specific stimuli, similar to Tamoxifen-inducible *Fos^Cre-ERT^*^2^ (e.g., TRAP) ^31,35,36^. All analyses described herein were conducted with young adults one week after exposure to the novel environment (Figure 1C). AAVDJ-DIO:APEX2-mGFP was injected into the CA3 and CA1 of *Fos^DD-Cre^* mice to label co-activated glutamatergic PNs in both areas. We focused exclusively on excitatory circuits; occasionally labeled GABAergic interneurons were omitted. The rationale for this strategy is further elaborated below and in the accompanying supplementary materials.

**Figure 1.**
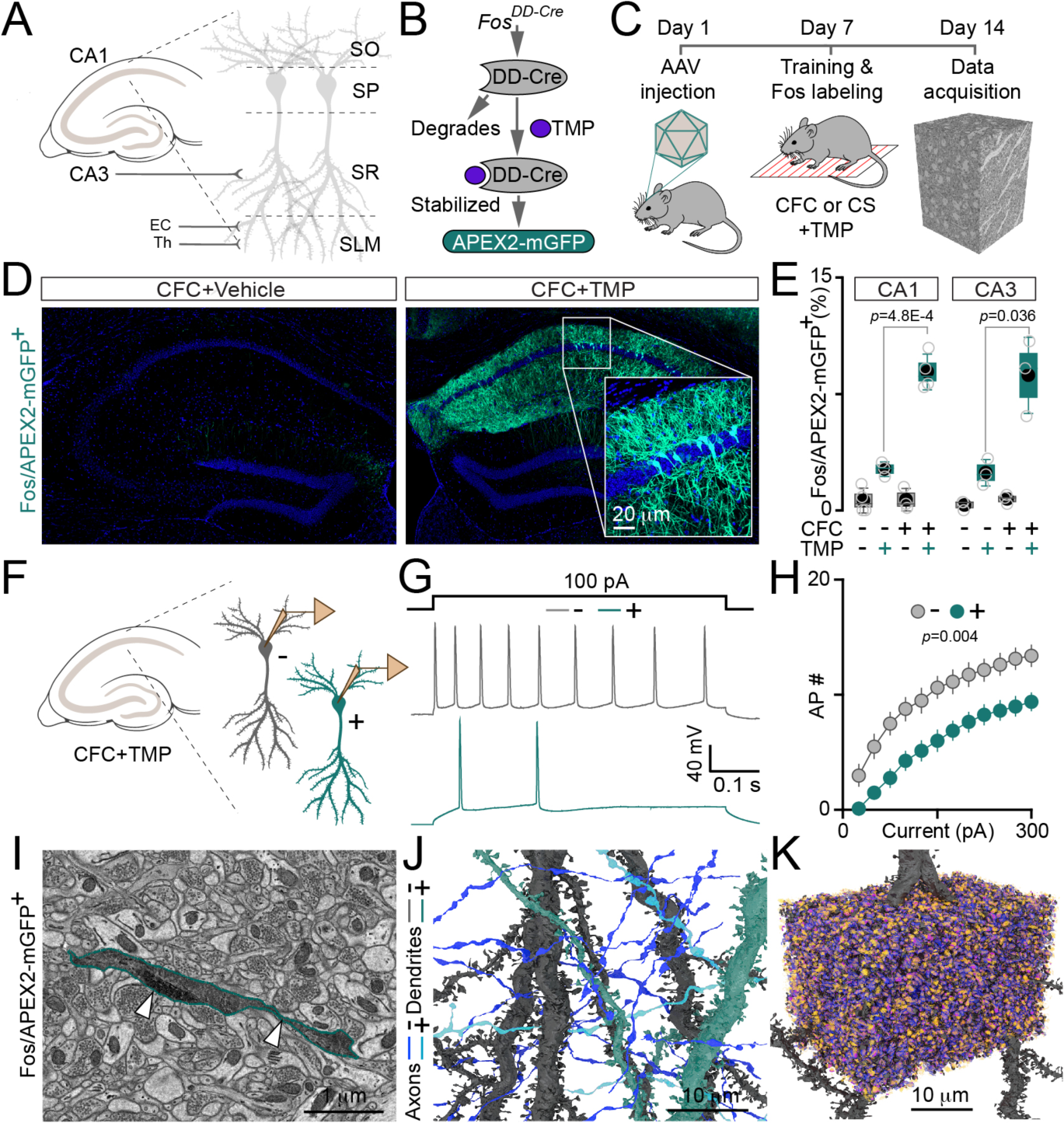
Workflow for identifying ultrastructural correlates of long-term memory in mouse hippocampus. **(A)** Schematics of excitatory pathways in the CA1. SO - stratum oriens; SP - pyramidal cell layer; SR - stratum radiatum; SLM - stratum lacunosum moleculare; EC - entorhinal cortex; Th - thalamus. **(B)** Irreversible Fos/TMP-dependent labeling of transiently activated neurons with APEX2-mGFP. See also Figure S1. **(C)** Overview and timeline of the experimental design. CFC - contextual fear conditioning; CS – neutral conditioned stimulus. **(D)** Confocal images of APEX2-mGFP fluorescence in the hippocampi of fear-conditioned *Fos^DD-Cre^* mice. Single doses of TMP (50 µg/g body weight) or a control vehicle solution were administered intraperitoneally 30 minutes post-training. **(E)** Comparison of background and TMP-induced APEX2-mGFP expression in trained and untrained animals. The quantifications of reporter-positive cells in areas CA1 and CA3 are shown. No CFC + vehicle, *n* = 4 mice; no CFC + TMP, *n* = 3; CFC + vehicle, *n* = 3; CFC+ TMP, *n* = 4. In this and subsequent similar panels, graphs display individual data points (open circles), mean values (filled circles), standard errors (boxes), standard deviations (vertical whiskers), and medians (horizontal lines). The *p* values were determined by t-test. **(F** to **H)** Electrical properties of Fos/APEX2-mGFP-negative (-) and positive (+) CA1 pyramidal neurons (PNs), as assessed 7 days after CFC and TMP treatment. See also Figure S2. **(F)** Schematics of side-by-side whole-cell recordings in acute hippocampal slices. **(G)** Sample traces of action potentials (APs) evoked by a 100 pA current injection. **(H)** Numbers of APs (Mean ± S.E.M.), plotted relative to stimulus intensity. *n* =3 mice/8 neurons per group. The *p* value was determined by t-test. **(I)** A typical example of sparse APEX2 labeling in the EM micrograph from fear conditioned mice (axonal fiber marked by white arrows with edges pseudocolored in green). **(J)** 3D views of PN dendrites and incoming Schaffer collateral (SchC) axons in the CA1sr of fear conditioned mice. APEX2 signal is pseudo colored as indicated in the legend. Only a few unlabeled projections are displayed. **(K)** Saturated reconstruction of excitatory synapses. The panel shows ∼1/12^th^ of the typical 3D image stack used for analysis. See also Figures S3 to S6.

We optimized the protocol for reliable induction of APEX2-mGFP with one TMP dose. The drug was acutely administered via peripheral intra-peritoneal (i.p.) injections 30 minutes post-training to allow DD-Cre transcription and translation. Confocal imaging of GFP fluorescence in hippocampal sections confirmed that the method works as designed. The expression of APEX2-mGFP was only observed after TMP delivery, and it was strongly upregulated by CFC. Compared to control TMP-treated mice, the fear-conditioned mice had a ∼4.5-fold increase in the quantities of APEX2-mGFP-positive neurons in pyramidal cell layers of CA1 and CA3 from ∼1.8% to ∼8% of the total populations (Figures 1D and 1E). These results agree with previous reports documenting the cellular expression of native *Fos* and other *Fos*-based fluorescent reporters in similar experimental settings ^2,8,9,12^. Moreover, our side-by-side whole-cell current-clamp recordings from labeled (+) and neighboring unlabeled (-) CA1 PNs in acute brain slices from fear-conditioned mice demonstrated that these neurons had markedly distinct electrical properties. The former had higher thresholds for action potential firing, likely reflecting a homeostatic dampening of intrinsic excitability to preclude the promiscuous reactivation by irrelevant stimuli when memory is already allocated (Figures 1F to 1H and S2A to S2C). In contrast, (+) PNs had no detectable widespread changes of excitatory synaptic strength, as evidenced by voltage-clamp recordings with protocols for the assessment of the probability of synchronous transmitter release from glutamatergic terminals and the amplitudes of evoked postsynaptic currents mediated by AMPA and NMDA receptors (Figures S2D to S2H).

### Reconstruction of excitatory circuits in the CA3-CA1 pathway

Having validated the chemogenetic toolbox, we sampled ∼100,000 μm³ SBEM stacks from the CA1sr of *Fos^DD-Cre^* mice that underwent two different procedures followed by an acute induction of APEX2-mGFP with TMP. All animals were placed in identical chambers, but only one group received foot shocks to distinguish the impacts of negative valence stimuli that prompt associative learning (CFC) and neutral conditioned stimuli (CS) alone. To ensure that fear-conditioned mice had retrievable memory 7 days later, at the time of tissue collection for SBEM, the behavior of separate cohorts from both groups was assessed upon their return to the original context (Figure S3). The 3D-EM stacks were aligned from raster images with 4 nm pixels, 2 μs dwell time, and 60 nm Z steps (Figures 1I and S4). Each stack contained ∼200,000 synapses. To expedite image processing, we built a pipeline for automatic segmentation of subcellular structures with an enhanced version of the previously described cloud-based convolutional neural network, CDeep3M ^37,38^. This deep learning AI platform was tailored for the retraining on manually segmented ground truth labels in Amazon Web Services (AWS) to generate accurate predictions of plasma membranes, axonal and dendritic shafts, nerve terminals, dendritic spines, postsynaptic densities (PSDs), and intracellular organelles (Figures 1J, 1K, S5, and S6). The precision of these predictions was verified by benchmarking against manual reconstructions from our own and previously published EM datasets ^28^. Excitatory circuits were reconstructed in automatically segmented images using the following criteria: 1) Each glutamatergic synapse with a characteristic PSD was assigned an unique ID and Euclidean coordinates; 2) Synapses, projections, and local connectomes were categorized by histories of activity on pre- and/or postsynaptic sides, as revealed by experience-dependent APEX2-mediated labeling of SchC axons and CA1 PN dendrites (Figure 1J); 3) Various morphological features of (-) and (+) PNs were compared within specimens and in different mice; 4) Because the (+) PNs were vastly outnumbered by (-) PNs (Figure 1E), we made a random selection of corresponding internal controls for all quantifications to balance the sample sizes and prevent heteroscedasticity; 5) When choosing data fitting and statistical methods, we took into account that distributions of structural synaptic parameters had asymmetrical Lognormal shapes ^39,40^.

### PNs with a remote history of activity during learning have unaltered distributions of individual synaptic sites and show no preference for wiring with each other

To begin elucidating how the architecture of the CA3-CA1 pathway is regulated by negative valence stimuli, we examined the reconstructions of CA1sr from mice exposed to CFC (Figure 2A). Initially, we focused on the dendrites of CA1 PNs and mapped their spines, postsynaptic compartments that cluster receptors and other signaling molecules near the sites of transmitter release from opposing axonal terminals (Figures 2B and S7). Sensory experience promotes the growth of new spines throughout the hippocampus ^3,41^. However, this dynamic growth appears to be counterbalanced by spine elimination ^3,41,42^ and its long-term consequences on neuronal wiring are uncertain. Our analysis showed that the spatial distributions and total spine counts on the dendritic arbors of (-) and (+) neurons were virtually indistinguishable (Figure 2C). We then mapped the presynaptic terminals along SchC axonal fibers coming from the CA3 (Figures 2D, S8A, and S8B). Given that the distances between neighboring terminals are more than 10-times greater than those between spines, we measured their lengths with an algorithm that accounts for the variability in axonal shaft curvature. Again, the distributions and total terminal counts on (-) and (+) axons were nearly identical (Figure 2E). To determine if task-specific co-activation of CA3 and CA1 PNs is attributed to their preference for wiring with each other, we compared the patterns of reporter expression in SchC fibers and their postsynaptic partners within local receptive fields (Figures 2F and S8C). The majority (∼85%) of terminals formed by (+) fibers contacted the spines of (-) dendrites, whereas the percentages of (+) spines receiving inputs from (-) and (+) axons were not significantly different (Figure 2G). In addition, (-) and (+) axons had similar branching and innervated the full spectrum of morphologically diverse spines, albeit the relative fractions of thin (T) and mushroom (M)-type spines were slightly uneven (Figures 2H, 2I, S8D and S8E).

**Figure 2.**
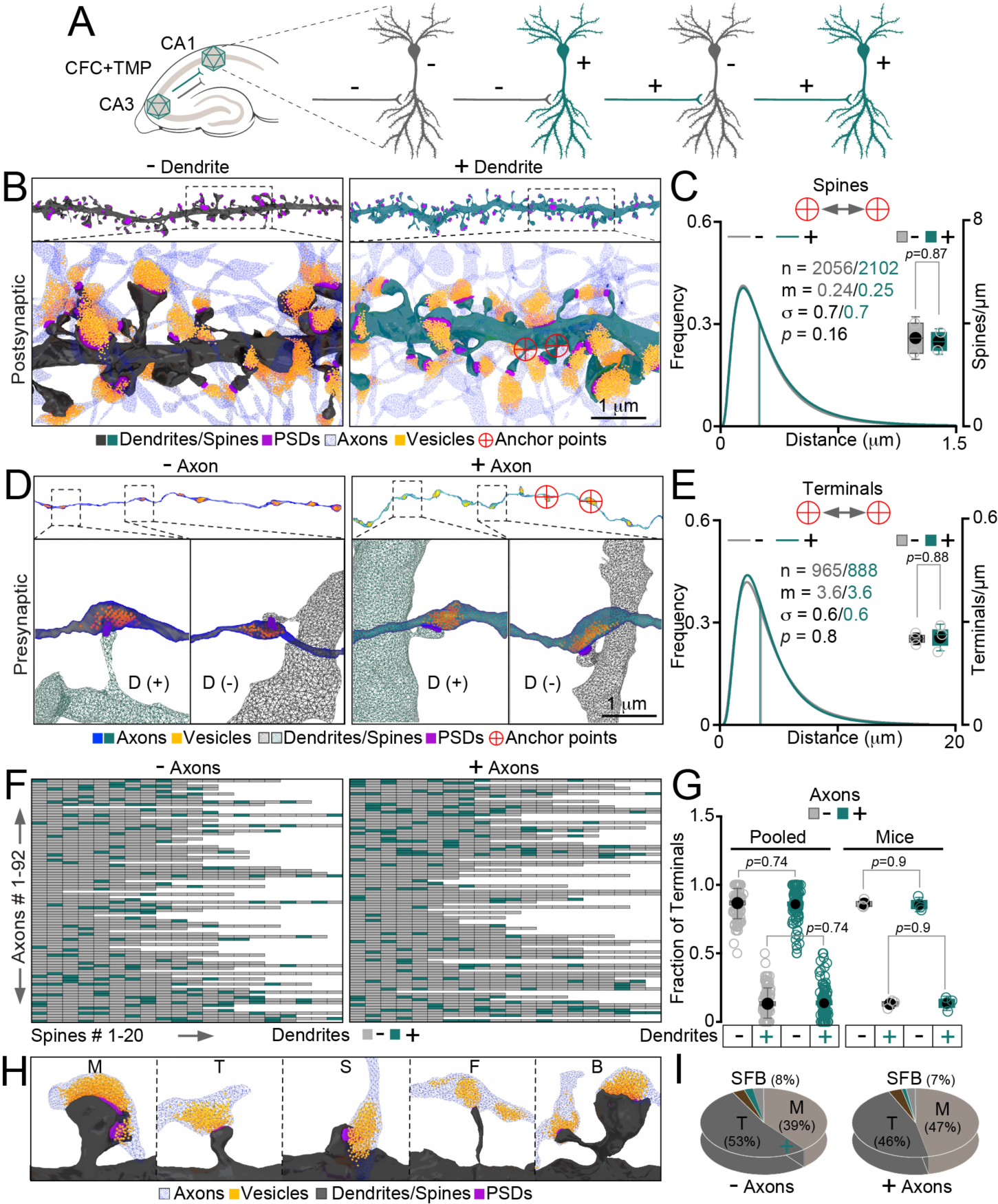
Local connectomes of PNs in the CA3-CA1 pathway. Excitatory synaptic networks were reconstructed in SBEM volumes from the CA1sr of fear-conditioned mice (7 days post-CFC). **(A)** Schematics of experience-dependent labeling and possible connectivity between Fos/APEX2-mGFP-negative (-) and positive (+) PNs. **(B)** 3D views of dendritic branches of (-) and (+) CA1 PNs. Enlarged images of regions marked by dashed boxes display local connectomes. Subcellular structures are colored as indicated in the legend. The anchor points exemplify the Euclidian coordinates of spines used for quantifications. Scale bar applies to both panels. See also Figure S7. **(C)** The combined graph shows the distributions of distances between spines (Lognormal curves with fitting parameters; in this and all subsequent similar panels, the sample sizes (n), medians (m) and standard deviations of logarithmic values (σ) are indicated) and the total spine counts per dendrite length, assessed across different mice (box with data overlap plot on the right). **(D)** 3D views of SchC axons of (-) and (+) CA3 PNs terminating in the CA1sr. Magnified images of areas marked by dashed boxes display pre- and postsynaptic structures. D - dendrites. Scale bar applies to all panels. **(E)** Distributions of distances between terminals along individual axons (Lognormal curves with fitting parameters) and the total terminal counts per fiber length (box with data overlap plots on the right). **(F)** Heatmaps representing the wiring of (-) and (+) axons with (-) and (+) dendrites. The color-coded boxes in vertical columns denote single spines innervated by different axonal fibers (horizontal lines) passing the receptive fields. See also Figure S8. **(G)** Fractions of presynaptic terminals formed onto the dendrites of (-) and (+) CA1 PNs. Both pooled datasets and values from individual mice are shown. **(H)** Examples of synapses with morphologically distinct spines. M - mushroom; T - thin; S - stubby; F - filopodia; B - bifurcated. **(I)** Fractions of M- T- S- F- and B-type spines innervated by (-) and (+) axons. All quantifications are from 3 separate mice. The *p* values were determined by Mann-Whitney tests and t-tests for distribution fits and box plots, respectively.

### Initial engram ensembles expand their connectomes through MSBs

While the bulk of *en passant* synapses of long-range glutamatergic axons have one-to-one spine-to-terminal ratios, some terminals are connected to multiple spines, usually residing on different dendritic arbors ^43–45^. Owing to their putative ability to synchronize outputs to several downstream targets, these atypical multi-synaptic boutons (MSBs) are thought to improve the fidelity of neurotransmitter signaling within circuits and coordinate the induction of Hebbian plasticity for sensory processing and learning ^45–48^. Yet, the precise functional roles of MSBs, the mechanisms of their morphogenesis, and their contribution to the assembly of cellular substrates of memory engrams are unclear. Since PNs with a remote history of activity correlated with fear learning had no changes in the overall repertoires of terminals and spines in the area CA1sr (Figures 2C and 2E), we hypothesized that the acquisition of new memories involves a shift in the equilibrium among conventional single-synaptic boutons (SSBs) and MSBs. To test this hypothesis, we identified MSBs in the CA3-CA1 pathway and studied their origins, arrangements on axons, structural characteristics, and synaptic target selectivity (Figure 3A). Although MSBs were present on both the (-) and (+) SchC axons of fear-conditioned mice, their counts on (+) axons were 1.6-fold higher. Consequently, (+) axons had a proportional loss of SSBs, and their total spine-to-terminal ratios were elevated by 18% (Figures 3B, 3C, S9A and S9B). Further analysis demonstrated that (+) MSBs were coupled with ∼50% larger fractions of all innervated spines than their (-) counterparts due to differences in both MSB abundance and their complexity (Figures 3D and 3E). Indeed, 26% of (+) MSBs contacted more than two spines, representing a ∼2.4-fold increase from the 11% in (-) controls. The maximum number of spines per MSB also rose from 4 to 6 (Figure 3F). These effects increased the sizes and diversity of connectomes of (+) axons, as demonstrated by quantifications of separate dendritic branches receiving inputs from each terminal (Figure 3G), fractions of compound synapses formed by SchC fibers onto neighboring spines on the same dendrite (Figure 3H), and averaged numbers of different dendrites whose spines are contacted by each MSB (Figure 3I). Remarkably, none of these alterations occurred in mice that were subjected to a neutral CS, indicating that axonal networks are reorganized in the CA1sr through MSBs in a stimulus-specific manner (Figures 3B to 3I). Hence, the restructuring of excitatory circuits observed after CFC is a signature of associative learning rather than the outcome of generic excitation of PNs or the *Fos*-driven expression of APEX2-mGFP per se. Nonetheless, the MSB target selection was not biased towards co-activated CA1 PNs. Consistent with our initial assessment of wiring diagrams (Figures 2F and 2G), ∼85% inputs from MSBs of (+) axons of fear-conditioned mice were provided to (-) dendrites (Figures 3J, 3K, S9C and S9D). Though the numbers of (-) and (+) MSBs on (+) dendrites were marginally mismatched (Figure 3L), this mismatch simply mirrored the unequal MSB availability.

**Figure 3.**
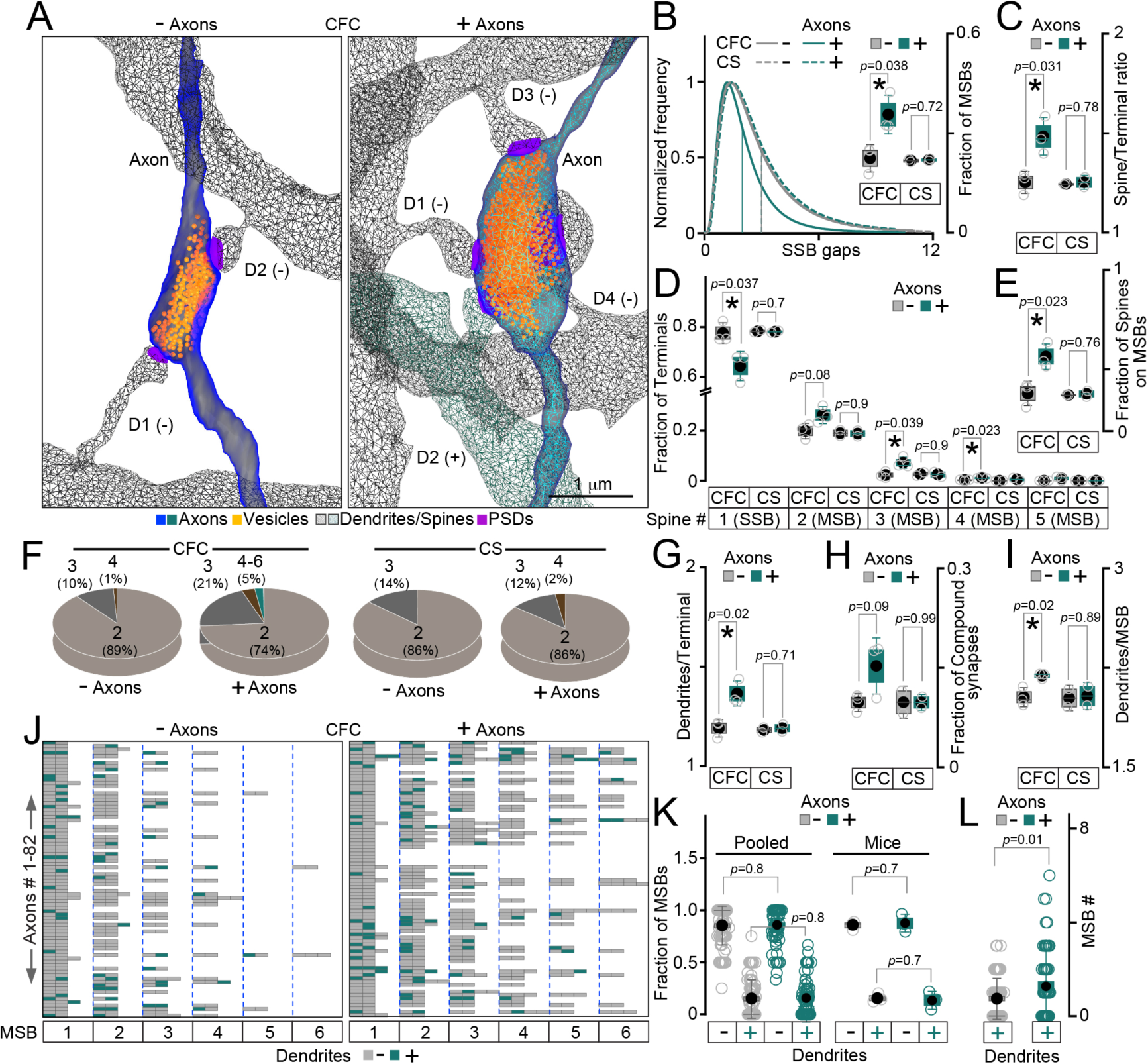
Stimulus-specific restructuring of excitatory circuits through MSBs. The spatial distributions, complexity, and connectivity of multi-synaptic boutons (MSBs) of SchC axons were examined 7 days after experience-dependent labeling of cellular ensembles in the CA3-CA1 pathway, as depicted in Figure 2A. **(A)** 3D views of MSBs of Fos/APEX2-mGFP-negative (-) and positive (+) axons connecting to distinct dendrites (marked as D1-D4) of (-) and (+) CA1 PNs in fear-conditioned mice. Scale bar applies to both panels. **(B** to **I)** Comparison of the long-term effects of presynaptic activity associated with fear learning (CFC, *n* = 3 mice) versus a neutral context exposure alone (CS, *n* = 2). Datasets were categorized based on axonal labeling. **(B)** The combined graph shows the count distributions of conventional single-synaptic boutons between MSBs (SSB gaps) and the averaged fractions of MSBs relative to SSBs on each axon, as determined in individual mice (box with data overlap plots on the right). **(C)** Overall spine-to-terminal ratios for (-) and (+) axons. **(D)** Fractions of terminals contacting indicated numbers of spines. See also Figure S9. **(E)** Fractions of spines contacted by MSBs of (-) and (+) axons. **(F)** Pie charts show percentages of MSBs innervating different numbers of spines, ranging from 2 to 6. **(G)** Numbers of dendrites innervated by individual terminals of (-) and (+) axons. **(H)** Fractions of compound synapses formed by (-) and (+) axons onto neighboring spines on the same dendritic branches. **(I)** Numbers of different dendrites innervated by MSBs of (-) and (+) axons (compound synapses were excluded). **(J** to **L)** Analysis of MSB networks in fear conditioned mice (*n* = 3). **(J)** Heatmaps representing axonal wiring via MSBs. Each box in the vertical columns shows the numbers of color-coded postsynaptic counterparts of each MSB formed by individual axons (horizontal rows, SSBs are omitted). See also Figure S9. **(K)** Fractions of MSBs of (-) and (+) axons connecting to (-) and (+) dendrites. Graphs demonstrate pooled datasets and measurements from different mice. **(L)** Cumulative counts of MSBs of (-) and (+) axons terminating onto (+) dendrites of previously activated PNs residing in the CA1. The *p* values were calculated by Mann-Whitney tests and t-tests for distribution fits and box plots, respectively.

### Memory encoding involves local input-specific restructuring of glutamatergic synapses

Thus far, our results suggest that the hippocampal PNs allocated for memory encoding expand their connectomes via MSBs while preserving the stable configurations of isolated synaptic sites along axonal and dendritic shafts. This expansion is driven by signals from presynaptic neurons regardless of the co-activation status of their downstream partners. The apparent recruitment of Fos-negative neurons that were not engaged in the immediate response to external cues is seemingly at odds with the Hebbian model. To explore this phenomenon, we further examined the impacts of negative valence and neutral stimuli on the structure of glutamatergic synapses (Figure 4A). First, we extrapolated synaptic weights by plotting the Lognormal distributions of spine head and nerve terminal volumes. These parameters are generally indicative of functional strength; for instance, spines become enlarged during long-term potentiation (LTP) and shrink during long-term depression (LTD) ^41,49^. In agreement with our electrophysiological recordings of evoked AMPA currents (Figure S2), (-) and (+) CA1 PNs of fear-conditioned mice had no detectable differences in spine head volumes, as assessed across dendritic branches of parent neurons irrespective of axonal labeling (Figure 4B). However, of two dispersed spine populations categorized solely by the activity history of contacting axons, the spines innervated by (+) SchC fibers were enlarged (Figure 4C). This enlargement was accompanied by decreased variability in spine sizes, as reflected by the lower standard deviation of logarithmic values (σ). The terminals of (+) axons were also markedly bigger, but their sizes became more variable (Figures S10A to S10C). In contrast, the volumes of post- and presynaptic compartments were unaltered in (+) PNs of mice subjected to CS (Figures 4D, 4E, S10D and S10E). To elucidate the discrepancies within the CFC datasets, we subdivided synapses by bouton type and repeated the distribution fits using axonal labeling as a frame of reference. The spines coupled to SSBs and MSBs of (+) SchC fibers were enlarged to similar degrees with 1.4- and 1.3-fold increases in median (m) head volumes, respectively. In each group, the σ’s were lower compared to corresponding (-) controls (Figures 4F and 4G). Likewise, (+) SSBs and (+) MSBs had the same 1.6-fold increase of median terminal volumes, despite the differences in bouton morphology. Even so, only (+) MSBs had much higher size variability than their (-) counterparts, as indicated by the σ value and the elongation of the distribution tail (Figures 4H and 4I). Conversely, no changes in the median sizes of either synapse type occurred in (+) PNs activated by CS alone (Figures S10F to S10I).

**Figure 4.**
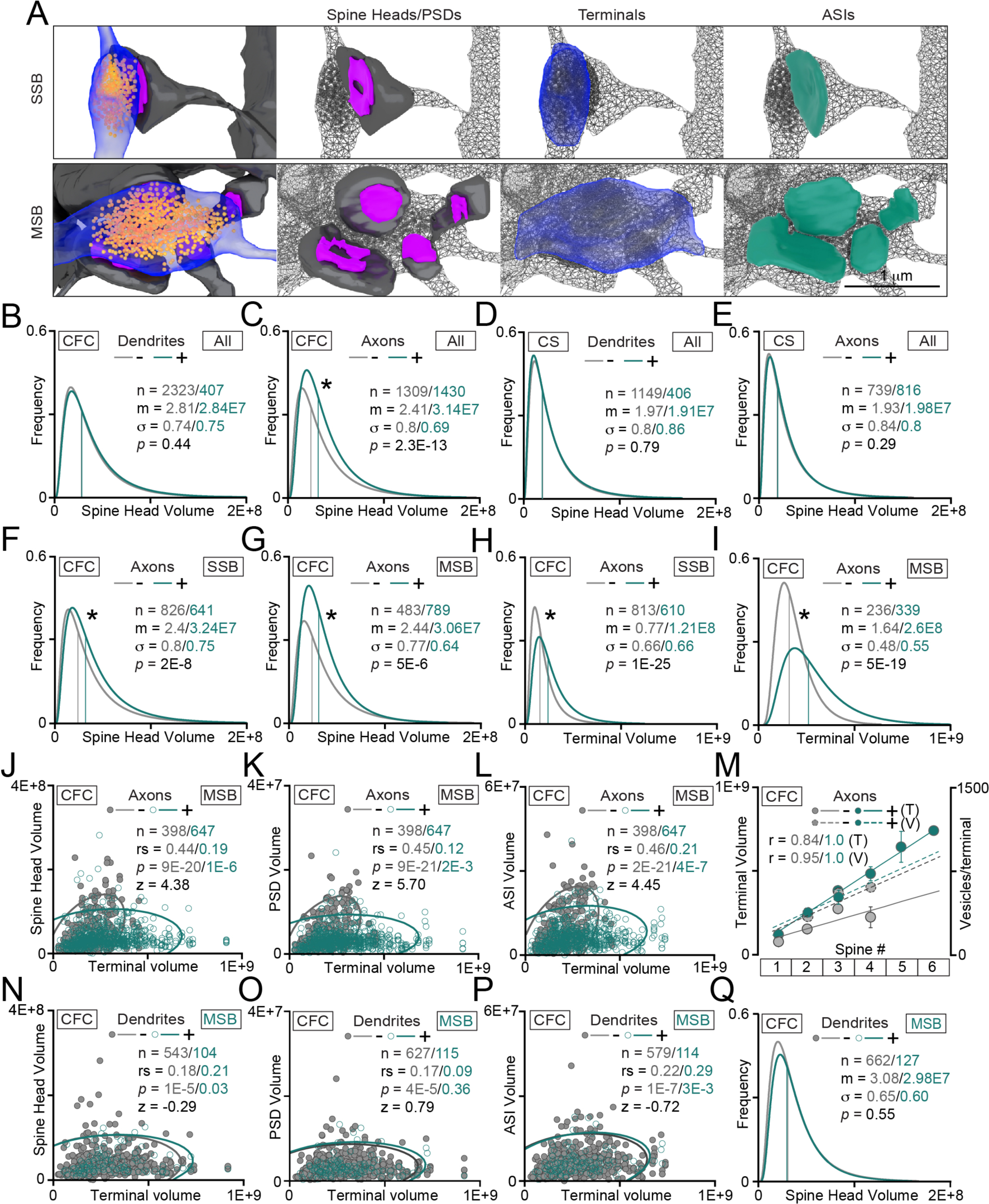
Stimulus-specific restructuring of individual excitatory synapses. Synapse sizes of PNs with remote histories of activity correlated with fear learning (*n* = 3 mice) or exposure to neutral CS (*n* = 2) were examined in the CA1sr using Fos/APEX2-mGFP labeling on post- or/and presynaptic sides as frames of reference. See also Figures S10 and S11. **(A)** 3D views of SSB- and MSB-type synapses with highlighted areas of spine heads, PSDs, axonal terminals, and axon-spine interphases (ASIs) used for quantifications. Scale bar applies to both panels. **(B)** Head volume distributions for all spines on dendrites of (-) and (+) CA1 PNs in fear-conditioned mice (datasets were categorized by postsynaptic label only). **(C)** Head volumes of spines innervated by (-) and (+) SchC fibers in fear-conditioned mice (presynaptic label only). (**D** and **E**) Same parameters as in panels (B) and (C) were assessed in mice subjected to CS without foot shocks. **(F** to **I)** Analyses of SSB- and MSB-type synapses in fear-conditioned mice, grouped by histories of CA3 neuron activity. **(F)** Head volumes of spines contacted by SSBs. **(G)** Head volumes of spines contacted by MSBs. **(H)** SSB terminal volumes. **(I)** MSB terminal volumes. **(J** to **L)** Correlations between indicated parameters in MSB-type synapses of fear-conditioned mice, grouped by histories of CA3 neuron activity. Scatter plots with confidence ellipses, sample sizes (n), Spearman correlation coefficients (rs), Fisher transformation scores (z) and *p* values are shown. **(J)** Spine head vs terminal volumes. **(K)** PSD vs terminal volumes. **(L)** ASI vs terminal volumes. **(M)** Combined plot shows the relationships between the terminal volumes (T) and vesicle pool sizes (V) vs numbers of innervated spines (1 spine - SSB; 2 or more spines - MSB). For each pairwise comparison, the Pearson correlation coefficients (r) are indicated. **(N** to **Q)** Individual postsynaptic targets of MSBs formed by (+) SchC axons were grouped into two categories based on expression of the reporter in CA1 PNs. Panels N to P show the correlations between the volumes of terminals vs spines, PSDs and ASIs. Panel Q shows the distributions of spine head volumes. All volumetric measurements are displayed in nm^3^.

Next, we calculated Spearman rank correlations to define the relationships between extrapolated post- and presynaptic weights in individual connections with unique IDs. Considering that the core components of typical excitatory synapses tend to be size proportional, we performed pairwise comparisons of volumetric measures for terminals, spines, PSDs and axon-spine interfaces (ASIs) (Figures 4A and S11A to S11D). While the magnitudes of input-specific effects on the median sizes of each of the opposing compartments appeared uniform in the SSB- and MSB-type synapses of fear conditioned mice at the population level, the structural proportionalities of single units were clearly distinct. The volumes of SSBs formed by (-) axons strongly correlated with those of contacted spines (rs=0.66), PSDs (rs=0.64) and ASIs (rs=0.64), whereas the correlations between the same parameters in (-) MSBs were only moderate (rs=0.44 to 0.46). These differences were even more pronounced among the (+) SSBs and (+) MSBs of previously activated CA3 PNs (rs=0.45 to 0.48 vs rs=0.12 to 0.21), though in both cases the correlations were weakened at the expense of asymmetrically enlarged terminals (Figures 4J to 4L and S11E to S11G). At the same time, the sizes of presynaptic boutons and the numbers of neurotransmitter vesicles in each bouton strongly correlated with the numbers of innervated spines (Pearson r=0.84 to 1.0; Figure 4M). The latter explains the physiological relevance of changes in (+) MSB morphologies, as discussed below. To explore the ramifications of learning-related remodeling of excitatory synapses in the context of the Hebbian rule, we then subdivided connections of (+) MSBs by the activity history of CA1 PNs. Under these experimental settings, neither the distributions of spine head volumes on (-) and (+) dendrites nor the correlations between pre- and postsynaptic structures were significantly different, indicating that the tuning of postsynaptic weights is not restricted to co-activated pairs (Figures 4N to 4Q). Taken together, these results support four conclusions: 1) Memory encoding coincides with sustained augmentation of synaptic weights, as evidenced by shifts in the distributions of synapse sizes elicited by CFC but not CS; 2) In line with our analyses of neuronal wiring, these events are prompted by presynaptic mechanisms that do not require activation of postsynaptic neurons above the threshold for *Fos* induction. Hence, the persistent re-structuring of CA3-CA1 synapses after associative learning appears to be non-Hebbian at a cellular level; 3) The experience-dependent growth of (+) MSB terminals, which leads to drastic loss of their structural proportionality with isolated spines, is attributed to higher complexity of target interactions. In other words, the asymmetric expansion of presynaptic boutons and vesicle pools fulfills the need to maintain neurotransmission with increased numbers of postsynaptic partners (compare Figures 3F and 4M); 4) Lastly, the observed changes in 0 values for spines innervated by (+) axons may reflect the matching of postsynaptic weights for optimal population coding.

### Presynaptic activity correlated with learning differentially affects the mitochondria and SER

To gain additional insights into the structural determinants of glutamatergic synapses of PNs activated during associative learning, we reconstructed membrane organelles essential for energy metabolism, protein synthesis, and intracellular calcium signaling - the mitochondria and smooth endoplasmic reticulum (SER) (Figures 5A, 5B, and S12A to S12C).

**Figure 5.**
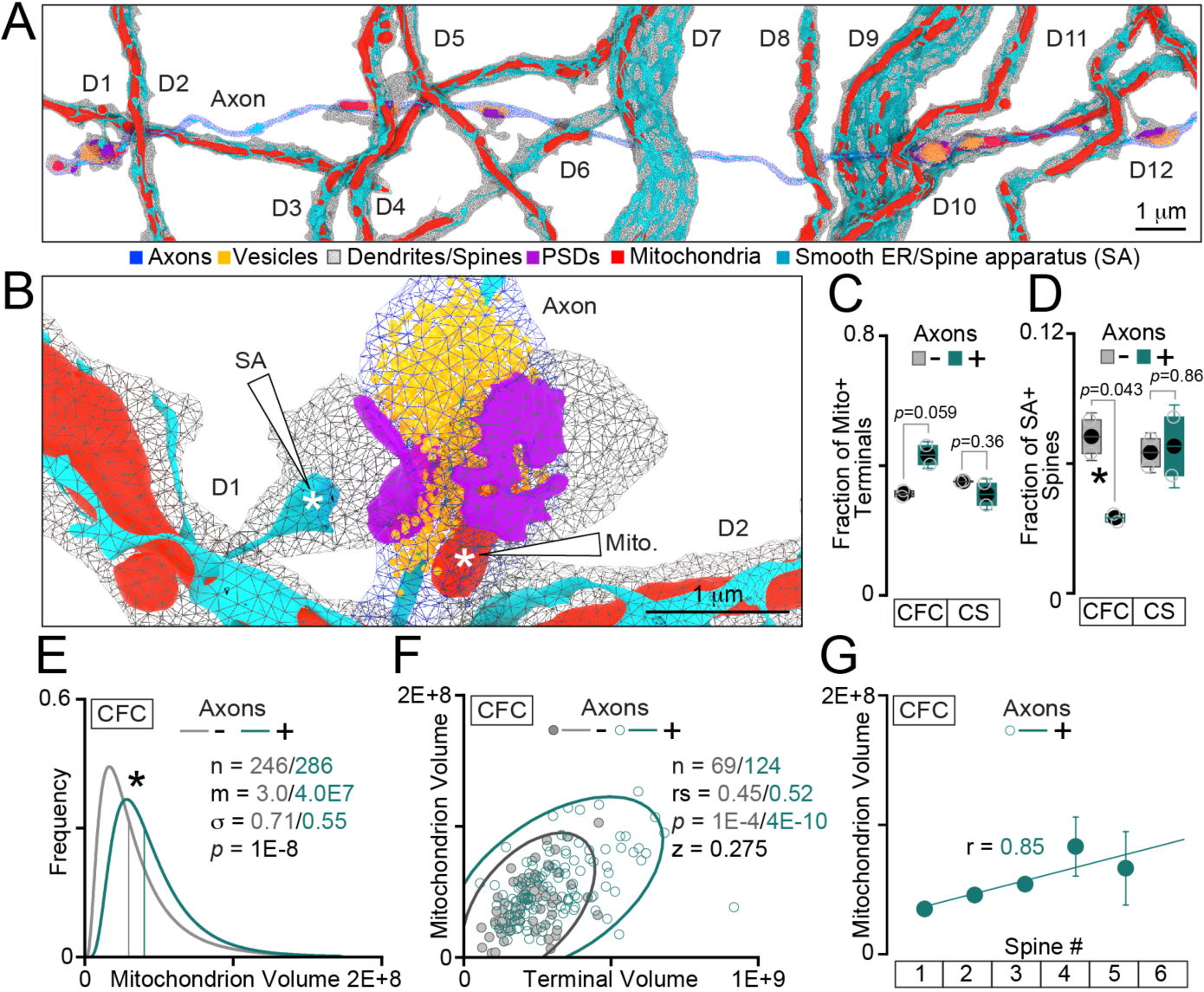
Effects of presynaptic activity correlated with learning on organelles residing in terminals and spines. **(A)** Typical 3D reconstructions of mitochondria and smooth endoplasmic reticulum (ER) in SchC axons and their target dendrites of PNs in the CA1sr. **(B)** Enlarged image with presynaptic mitochondria and postsynaptic spine apparatus (SA) marked by arrows and asterisks. **(C** to **G)** Analyses of mitochondria and SA in synapses formed by Fos/APEX2-mGFP-negative (-) and positive (+) axons (all datasets were categorized by the presynaptic label). **(C)** Fractions of axonal terminals containing mitochondria in mice subjected to CFC or CS alone. **(D)** Fractions of spines containing SA. See also Figure S12. **(E)** Volumes of mitochondria anchored in (-) and (+) terminals of fear-conditioned mice. Lognormal distribution curves and fitting parameters are shown. **(F)** Relationships between mitochondrion and terminal volumes. Scatter plots display confidence ellipses, sample sizes (n), Spearman correlation coefficients (rs), Fisher transformation scores (z) and *p* values. **(G)** Correlation between the volumes of presynaptic mitochondria and the numbers of spines innervated by each mitochondria-containing terminal of (+) axons (1 spine - SSB; 2 or more spines - MSB). All quantifications are from 2 mice/group. The volumetric measures are displayed in nm^3^. The *p* values were calculated by t-tests and Mann-Whitney tests for box plots and distribution fits, respectively.

In addition to supplying ATP for SNARE-mediated vesicle fusion and recycling, the mitochondria anchored at presynaptic boutons modulate neurotransmitter release kinetics by preventing calcium buildup triggered by repetitive action potentials ^50–52^. On the other hand, the mitochondria spanning the dendritic shafts seldom enter the spines ^21^. The postsynaptic calcium homeostasis of glutamatergic PNs as well as local translation in spines are thought to be regulated by a part of the SER network called the spine apparatus (SA) ^53,54^. Curiously, many excitatory synapses lack the mitochondria and SA ^38,55^, but the underlying reasons and the functional implications of this heterogeneity are poorly understood. Prior EM studies, including recent work from our group, raise the possibility that subcellular localization of these organelles is influenced by sensory stimuli. However, this notion is based on indirect observations in electrically stimulated brain slices and excitatory neurons whose glutamate secretion was permanently blocked *in vivo* throughout development ^38,55^. We found that, regardless of sensory experience and neuronal activity source, the mitochondria were more likely to be anchored in MSBs than in SSBs. The fear conditioned mice tended to have more mitochondria-containing terminals along (+) SchC axons, although this difference was not statistically significant (Figures 5C and S12D). On the contrary, the spines receiving inputs from (+) SchC axons of CA3 PNs had a notable CFC-specific loss of SA regardless of presynaptic bouton type (Figures 5D and S12E). Because efficient calcium buffering is necessary for the synchronization of vesicular release and this process depends on both the anchoring of axonal mitochondria and their size ^56,57^, we asked whether these organelles undergo compensatory expansion in enlarged terminals of PNs activated by CFC to meet the higher energy demand and limit the diffusion of calcium ions upon entry from the extracellular space. Such homeostatic mechanism could be particularly important for MSBs in which vesicles from a shared pool undergo exocytosis at multiple sites. Supporting this prediction, our quantifications demonstrated that the median mitochondrion volumes were increased in terminals of (+) SchC axons by 1.3-fold (Figure 5E). Moreover, the pairwise correlation analyses showed that the sizes of presynaptic mitochondria were proportional to both, the terminal sizes (rs=0.45 to 0.52) and the numbers of contacted spines (r=0.85) (Figures 5F and 5G). Hence, presynaptic activity correlated with learning differentially affects the organelles that maintain synaptic homeostasis and define the propensities for short- and long-term plasticity, by promoting the local function of axonal mitochondria and by disabling the SA in spines.

## DISCUSSION

In summary, our study reveals mechanisms that regulate the synaptic architectures of memory engrams, adhering to three core principles: i) their induction is driven by presynaptic activity associated with specific sensory stimuli; ii) their expression transcends simultaneously activated neurons, as marked by *Fos*; and iii) they involve multi-synaptic boutons (MSBs). The stimulus-specific remodeling of MSB networks and individual synapses may underlie representational drifts recently observed in the hippocampus and cerebral cortex through *in vivo* calcium imaging and electrophysiology ^14–19^. While our observations challenge the general applicability of the Hebbian model, they do not seek to refute it entirely. Instead, we argue that the Hebbian plasticity does not account for the wiring and sustained experience-dependent rewiring of the cellular substrates underpinning memory traces. It is noteworthy that prior optical imaging studies, which suggested hippocampal neurons enhance connectivity in a Hebbian fashion during fear learning, also utilized the *Fos* promoter to identify transiently activated neurons ^12,13^. Thus, the discrepancies in our findings might stem from differences in image resolution and/or the depth of circuit analysis, rather than inherited limitations of *Fos*-based expression systems.

The discoveries of experience-dependent synaptogenesis and synapse elimination have popularized the notion that sensory cues have lasting impacts on the computational attributes of central neurons by altering the number and/or subcellular distribution of their synaptic contacts ^1,3,41^. Our results highlight an alternative strategy whereby neurons engaged in memory encoding diversify the postsynaptic recipients of chemical signals from the same pools of neurotransmitter vesicles residing in each MSB, while maintaining the numbers of individual nerve terminals and dendritic spines in a steady state. This strategy is advantageous in adult brain regions where densely packed projections, synapses, non-neuronal cells, and components of the extracellular matrix pose topological barriers to the restructuring of axonal/dendritic arbors and synapse growth. It is intriguing to predict that MSB networks progressively change in the brain as more information is acquired over the lifespan and that MSB dysfunctions contribute to cognitive decline. Furthermore, the morphological features and physiological properties of these unusual synapses may reflect general intelligence both across and within mammalian species. This possibility is indirectly supported by preprinted 3D-EM analysis of a single specimen from the human cortex, indicating a higher density of MSB-innervated spines in humans compared to mice (https://doi.org/10.1101/2021.05.29.446289).

Although MSBs have been observed by EM across various brain regions ^44,45,48^, their molecular composition is yet to be unraveled. The patterning of MSBs along axons, their postsynaptic targets, and the intricate nature of these interactions are likely controlled by unique sets of surface adhesion, scaffolding, and signaling molecules ^58^. Given that MSBs are surrounded by conventional synapses on the same axon, identifying these molecules will necessitate the development of novel methodologies to isolate structurally distinct synapse sub-populations with shared cellular origins and neurotransmitter identities. Exploring the MSB composition will lay a roadmap for optical imaging and targeted manipulations of MSBs *in vivo*. Combined with behavioral readouts and mathematical modeling, these future studies will advance our understanding of the core principles of information processing in the brain and, possibly, will facilitate the design of next-generation algorithms for artificial intelligence.

## ACKNOWLEDGEMENTS

We thank Dr. Ardem Patapoutian and the members of the Maximov and Ellisman laboratories for advice and critical comments.

## Author contributions

MU, YZ, AM and ME conceived the study. YZ characterized the mouse models by optical imaging and prepared specimens for SBEM. FP performed electrophysiological recordings from acute brain slices. EB and CK collected SBEM stacks. MU and MH developed computational tools for image segmentation. MU oversaw all 3D reconstructions, performed behavioral experiments, and analyzed the EM data. KC, DL, LC, TQ, EZ, LP, EBeu, WZ, BD and EM assisted MU with reconstructions. AM wrote the manuscript.

## Data and materials availability

All reagents and resources, including expression vectors, mice, and algorithms for image analysis can be obtained from A.M. upon request.

**Supplementary materials** are enclosed with the manuscript and include materials and methods and 12 figures with legends.

## Supplementary Materials

## Materials and Methods

### Mouse lines and expression vectors

The *Fos^DD-Cre^* and *Camk2^Cre^* strains used in this study have been described previously ^31,59^. Mice were housed, crossed, and analyzed according to protocols approved by the IACUC committee. All animals were a mix of C57BL/6 and 129/SV backgrounds. Males and females were examined together. The APEX2-GFP and Cherry-H2B coding sequences were inserted into a well-characterized shuttle vector containing the EF1α promoter and the DIO cassette for inducibility with Cre ^59–61^. For constitutive expression of GFP, the coding sequence was subcloned into a similar vector containing the pan-neuronal Synapsin (Syn) promoter.

### AAV production and injections

AAVs were generated in house in HEK293T cells, purified by Heparin-based affinity chromatography, tittered by real-time quantitative PCR, and injected into brains of mice carrying the Cre drivers at titers of 2x10^12^ GC/ml as we have previously described ^59–61^. Mice were anesthetized with 1.5%– 2% isoflurane in O_2_; stereotaxic injections were performed bilaterally into the dorsal areas CA3 and CA1. Viruses were infused for 5 minutes at a rate of 100 nl per minute.

### Drug delivery

TMP-lactate (Sigma, Cat #T0667) was reconstituted in PBS prior to each experiment (30 mg/ml) and administered to mice by intraperitoneal injections through a 29g needle at a dose of 300 μg/gm body weight.

### Fluorescent imaging

Mice were anesthetized with isoflurane and perfused transcardially with 25 ml of ice-cold PBS followed by 25 ml of 4% PFA in PBS using a peristaltic pump. The brains were removed, incubated overnight in 0.4% PFA, and sliced in ice-cold PBS using a vibratome. 90 μm thick DAPI-stained coronal sections were imaged under the Nikon C2 microscope with 0.3-0.5 μm Z-intervals using 20x and 40x objectives. Digital manipulations were uniformly applied to all images.

### Behavior

Mice were allowed to explore the fear-conditioning boxes (Context A, Med Associates SD) for 3 minutes and were then subjected to four bursts of foot shocks (0.55 mA, 1 minute inter-shock intervals). The control groups were exposed to the same conditional visual and olfactory stimuli without foot shocks. Memory retrieval was tested 7 days later upon exposure to the original context. Freezing was measured for 3 minute intervals with 0.75 second bouts.

### Electrophysiology

Mice were anesthetized with isoflurane. Brains were removed and placed into an ice-cold oxygenated buffer (95%O2/5%CO2) containing 228 mM sucrose, 2.5 mM KCl, 0.5 mM CaCl_2_, 7 mM MgCl_2_, 26 mM NaHCO_3_, 1 mM NaH_2_PO_4_, and 11 mM glucose. Transverse, 300 μm thick slices were sectioned with a vibratome and initially stored at 32°C in oxygenated aCSF containing 119 mM NaCl, 2.5 mM KCl, 1 mM NaH_2_PO_4_, 26 mM NaHCO_3_, 1.3 mM MgCl_2_, 2.5 mM CaCl_2_, 11 mM glucose (pH 7.4, 292 mOsm), and then allowed to recover for 1 hour in oxygenated ACSF at 24 °C prior to recording. Membrane and synaptic currents were monitored in whole-cell current- and voltage-clamp modes, respectively, using Multiclamp 700B amplifier (Molecular Devices, Inc.). Recordings were performed at room temperature. The pipette solution contained 135 mM CsMeSO_4_, 8 mM CsCl, 0.25 mM EGTA, 10 mM HEPES, 2 mM MgATP, 0.3 mM Na_2_GTP, 5 mM QX-314, and 7 mM Na_2_phosphocreatine (pH 7.4, 302 mOsm). Synaptic responses were evoked by 1 ms local field stimulation using an extracellular electrode. Data were sampled and analyzed with pClamp10 (Molecular Devices, Inc.) and OriginPro (Origin Lab) software packages.

### Sample preparation for EM

Mice were anesthetized via intraperitoneal injections of ketamine/xylazine, transcardially perfused with oxygenated Ringer’s solution, and then perfused with a buffer containing 2% paraformaldehyde, 2.5% glutaraldehyde, 150 mM cacodylate and 2 mM CaCl_2_. The brains were post-fixed overnight in the same solution at 4 °C. 100 μm thick coronal slices were cut in Vibratome and prepared for SBEM imaging using the following sequential procedures: 1) Overnight post-fixation at 4 °C followed by washes in the buffer containing 150 mM cacodylate and 0.2 mM CaCl_2_; 2) Fixation at room temperature for 1 hour in 2% OsO_4_ in cacodylate; 3) Staining in 0.5% aq. thiocarbohydrazide; 4) Staining with 2% aq. OsO_4_; 5) Overnight incubation at 4 °C in 2% aq. uranyl acetate; 6) Staining with lead aspartate at 60 °C for 30 min; 7) Dehydration on ice in 70%, 90%, and 100% ethanol followed by dry acetone; 8) Infiltration with acetone:Durcupan ACM; 9) Embedding in 100% Durcupan resin at 60 °C for 48 hours. Approximately 1 mm square pieces of tissue were mounted on Gatan SBEM specimen pins with conductive silver epoxy. For steps 2 to 6, each procedure included subsequent washes in water at room temperature.

### Acquisition and analysis of 3D-EM stacks

#### SBEM imaging

Samples were imaged under the Zeiss Merlin scanning electron microscope equipped with a Gatan 3View. Imaging was performed at 2.5 kV and 85 pA using a focal charge compensation device to minimize specimen charging (2.5x10^-3^ mbar nitrogen gas). ∼110,000 μm^3^ stacks were collected from dorsal CA1sr using 15k x 15k raster images with 4 nm pixels, 2 msec pixel dwell time, and 60 nm Z steps. Acquired volumes were aligned in IMOD ^62^.

#### Deep learning-based image segmentation

Automatic segmentation of different subcellular structures was performed with version two of CDeep3M, an image segmentation platform that utilizes a convolutional neural network (CNN). This platform enables effective processing of multiple microscopy modalities, including SBEM. Network training and inference were accomplished using the Docker-based version of CDeep3M on both local lab computer GPUs and on the Amazon Web Service (AWS). Since training the network from scratch is time consuming, available pre-trained neural networks were retrained on the specific SBEM image sets utilized in this study. This method of domain adaptation reduces effort and time by 90% while still achieving high segmentation accuracy ^37^. Pre-trained models for plasma membranes, mitochondria, vesicles, and synapses were downloaded from the publicly available Cell Image Library and retrained for each image stack using small volumes with manually segmented ground truth labels. New models for dendritic spine heads and axon terminals were generated by training the network from baseline. All image contrasting and manipulations were done with Fiji/ImageJ ^63^. The accuracy of retrained neural networks was validated with small volumes from separate areas of the SBEM block. Validated models were then applied to automatically segment entire volumes. The output prediction maps were stitched together and used for either fully automated reconstruction in IMOD or semi-automated reconstructions in VAST ^21^.

For modeling the gross anatomy of neurites in the block, instance segmentation was performed using PyTorch Connectomics, a recently developed deep learning framework for automatic and semi-automatic annotation of connectomics datasets ^64^. The CNN was trained with small image stack in which all neurites and astrocytes were segmented. The trained network was used to predict an affinity map and a watershed algorithm was used to generate the dense segmentation. The automatic reconstructions from this pipeline were then used for 3D visualization.

#### Automatic reconstructions

Automated reconstructions were performed using IMOD and PyIMOD, a python library for manipulating IMOD files (https://github.com/CRBS/PyIMOD). Prediction maps of dendritic spine heads, axon terminals, synapses, vesicles, and mitochondria were initially generated in CDeep3M. Since each SBEM image set had slight variations in XY size, cropped versions of each prediction map were created with dimensions of ∼10,000 x 10,000 pixels x 500 Z-slices (48,000 um^3^). Vesicle analysis was restricted to a smaller volume (∼12,000 um^3^) because of heavy computational load. 2D contours of each feature were made by thresholding using *imodauto* and small contour artifacts were automatically removed by defining a minimum area. 3D meshes of each object (excluding vesicles) were then created using *imodmesh* and sorted into separated objects using *imodsortsurf*. PyIMOD was used for final post-processing by removing clipped objects touching the image set border using *removeBorderObjects* and filtered by contour number by designating a minimum and maximum slice number using *filterByNContours.* Object volumes and counts per block volume were extracted using *imodinfo*.

To obtain vesicle counts per terminal, the generated vesicle contour and final sorted axon terminal files were converted into TIFF format and imported into VAST with “Import Segmentation from Images” function. The individual vesicle coordinates were extracted with the Vasttools “Export Particle Clouds” function. Since the 60 nm z-steps of image stacks were larger than the vesicle diameters (∼40 nm), each separated 2D contour in each slice was considered a unique vesicle. A custom MATLAB script was written to count the number of vesicles in each terminal based on whether the coordinates of each vesicle contour overlapped with a segmented terminal of a specific ID.

Although there is a slight chance for merge errors to occur across closely located objects using this automated segmentation approach, we reasoned that this did not bias the results because the same thresholding and filtering criteria were applied to all experimental conditions. The absolute accuracy of our pipeline was further validated by comparing the small-scale automatic reconstructions with manual and instance-based segmentations of the same sample. We found that only few features were merged; importantly, there was no bias in merge errors across different samples.

#### Semi-automatic segmentation

Semi-automatic segmentations were carried out in VAST Lite version 1.4.1, which is a voxel painting program designed for the analysis of large volumetric datasets ^65^. We used masked painting in VAST in combination with the CDeep3M predicted boundary maps to constrain painted areas so that the outline of each object is traced automatically. By selectively painting neuropil structures, we were able to achieve quick and accurate reconstructions of axons, dendrites, synapses and mitochondria. Errors in painting from occasionally inaccurately predicted boundaries were manually corrected. For the segmentation of dendrites and axons, we used membrane prediction boundary maps for filling the outlines of each cellular structure. For the segmentation of mitochondria, we used a prediction map for masked painting to fill the entirety of each structure. Since the present study was focused on excitatory circuits, analyses were restricted to spiny apical dendritic segments of PNs and axons that form glutamatergic synapses with characteristic postsynaptic densities (PSDs). All segmented volume data were extracted in MATLAB through the included software VastTools.

#### Skeletonization and assignment of coordinates

Skeletons for traced dendritic and axonal segments were generated in VAST. Using the annotation function, connected nodes were placed along the center of each reconstructed structure’s cross-section from the first slice of the image volume to the last at spaced intervals. Image stacks where dendritic spines or axonal terminals were present along the skeletonized length were also marked in order to assess spine and terminal distributions and distances across the entire structure. Nodes were placed in inter-terminal spaces to account for the excess curvature seen in axons. In a few instances wherein the axon split and formed a new branch, a single path was reconstructed to maintain compatibility with most of the analyzed dataset. Annotated length data was extracted in MATLAB through the Vasttools functions and API.

#### Reporter tracing

APEX2-positive axons and dendrites were identified by the presence of dark peroxidase staining throughout their membranes and intracellular structures. A clear indicator of reporter expression was the increased intensity of staining in smaller compartments. For example, the staining in dendritic spines was much stronger than in shafts. Since the expression of the reporter was not cell-type specific, we also observed labeling of putative inhibitory neuronal structures that formed symmetrical, shaft-targeted synapses. While intriguing, these reconstructions were excluded from the final quantifications. The staining intensity of axonal fibers was notably darker than that of dendrites, and in certain cases, occluded the visibility of intracellular contents. Since axonal staining did not affect our ability to assess the distributions of individual synapses, all reconstructed excitatory axons were included. For the analysis of mitochondrial localization in terminals, a few axons were omitted in which the staining was too dark for segmentation. All blocks had comparable ranges of staining intensity and there was no difference in the fraction of excluded axons.

While we trained the convolutional neural network models on the image sets from this study, there were still a disproportionate amount of prediction errors for labeled structures due to the dark staining. This was particularly problematic in axons. In cases where prediction mistakes occurred, they were manually corrected or traced. It is possible that slight differences in volume measurements between labeled and unlabeled structures emerged from intrinsic variation amongst segmentation methods. However, virtually all documented volume changes of APEX2-positive structures of fear conditioned mice were not found in mice subjected to neutral CS, despite the same staining patterns. Hence, the possible differences in absolute volumes could not significantly affect the overall results.

#### Reconstructions of APEX-positive and -negative projections

We randomly selected and reconstructed sets of labeled and adjacent unlabeled dendrites with similar orientations. Although large, apical dendritic trunks were present in the CA1sr, we focused on smaller higher order branches that formed the majority of spines. The shaft diameters of selected pairs were always matched; there was only minimal variation between the entire sets. To further reduce the variability, spine measurements were carried out with shorter segments (∼100 spines/branch). Axons were chosen using the same criteria and reconstructed throughout the block. Some spine-innervating axons formed at least one synapse with a smooth, non-spiny dendrite. Additionally, there were occasional synapses onto dendritic shafts. These synapses were included in the measurements of synapse distribution along axonal shafts, but were omitted for volumetric measurements.

#### Classification of Terminals and Spines

Nerve terminals were classified based on the number of innervated spines as SSBs (1 spine) or MSBs (2+ spines). Because presynaptic boutons are usually separated by large distances across an axon, there was typically little ambiguity as to whether a spine belonged to one bouton or another. Furthermore, the pre- and postsynaptic partners were indicated by the visible PSDs. In the few cases where the MSB or SSB designation was unclear due to irregular shapes, the continuity of the vesicle pool was used to distinguish the terminal borders. Dendritic spines were classified into 5 morphologically distinct types: Mushroom (M), Thin (T), Stubby (S), Filopodia (F), and Bifurcated (B) as described previously ^66^. Rare spines whose shapes were unclear were excluded from quantifications except for measurements of distances between synapses.

#### Splitting of spine heads and terminals

Spine heads and nerve terminals were manually separated from their parent structures using the splitting tool in VAST. This procedure created new segments, which could then be automatically volume-filled with the filling tool. Spine heads were split at the base of the neck. For the analysis of synapses along projections, there were a few instances in which a synapse was formed onto non-spiny, smooth dendrites and dendritic shafts. This made up a small fraction of the total pool and was not included in the measurements of volumes. Terminals were split at each end where axonal cross-sections minimize, vesicle pools end, and/or opposing PSDs end.

#### PSD and ASI measurements

PSDs were identified as darkly stained regions at the ends of dendritic spines, typically opposing axonal terminals with synaptic vesicles. PSDs were segmented in VAST by adjusting the pen size depending on PSD thickness. Since there can be some variation in the visibility of the PSD depending on the spine orientation (for example, cross-sectioned synapses may seem more continuous than oblique synapses), we may have underestimated PSD volumes in a few cases. However, we reasoned that this variability did not significantly bias the results because we applied the same segmentation criteria to all conditions and analyzed large sample sizes. ASIs were defined as the entire contact area between the axon terminal and the postsynaptic spine. The ASIs were segmented in VAST with a fixed pen size across each plane of the synapse, except for oblique synapses wherein the entire contact zone was filled in the plane. Similar to PSD measurements, we used the same justification for possible underestimation of ASI measurements based on the synapse orientation.

#### Mitochondria and SA

Mitochondria-positive terminals were identified by the presence of one or more mitochondria within the boundaries of the bouton using the criteria described above. Mitochondria were semi-automatically segmented in VAST using masked painting with a CDeep3M mitochondria prediction map. In cases where the prediction map contained errors, the mitochondria were manually corrected or traced. Although the APEX2 reporter darkly stained presynaptic compartments, the boundaries of mitochondria were still clearly distinguishable. A few reporter-positive axons in which all intracellular contents were completely masked by APEX2 staining were excluded from analysis. Spines were defined as SA-positive when stacks of SER folds were visible in necks and/or heads. This population was differentiated from spines with a single tubular-SER.

#### Data extraction and analysis

Large-scale automated reconstructions modeled in IMOD were extracted in a Linux terminal using *imodinfo* and placed in .csv files for analysis. This method was used for the cubic density measurements of spine heads, terminals, mitochondria, and vesicles. All other data extractions were performed in VastTools, a MATLAB script that interfaces with VAST via included API. For example, VastTools was employed for extractions of volume measurements, coordinates for length and distance measurements, and 3D surface meshes. For all data extractions and 3D model exports, the same parameters were uniformly applied to all samples. Since VastTools permits quantifications at lower resolutions, we used the native image stack voxel size (4 nm × 4 nm × 60 nm) at Mipmap level 0. All numerical values were extracted before 3D modeling.

#### Volume measurements and 3D model extraction

Volumes of dendritic spines, axonal terminals, PSDs, ASIs, and mitochondria were measured with the Measure Segment Volumes function, which counts the total number of voxels of different specified objects in a boundary area. 3D models were generated with “Export 3D Models” function, which creates surface meshes in VAST and exports them as .obj files for modeling in Blender. Some features were exported as lower-resolution models (Mip 2-3) to make post-export smoothing easier. The Export Particle Clouds function was used to export 3D surface mesh models (.obj) of vesicles for 3D modeling in Blender.

#### Coordinate exporting

Whole skeleton length measurements were extracted using VastTools function “Measure Skeleton Lengths” to sum the overall distances of placed nodes along neurites. However, internode length measurement functions are not currently implemented in the VastTools window of the current version of VAST. Thus, we wrote our own MATLAB scripts that utilized API functions to extract these measurements. As stated above, we employed the *Annotation* tool in VAST to create skeletons nodes that were placed along neuropil structures across the image stack (e.g., axons). In order to prevent overestimation of length, nodes were placed in the center of each structure. To analyze dendrites, nodes were placed at the starts and ends of dendritic fragments, as well as in each section where a spine branched off from a shaft. This allowed us to extract parameters such as dendritic fragment length, linear spine density, and distances between individual spines. Every dendritic protrusion was considered a spine. While the skeletonization approach used for dendrites generally applies for axonal fibers, we wanted to account for higher curvatures of these structures. Hence, the nodes were placed at the beginning and end of each axon, in terminals with opposed spines, as well as in arbitrary sections along fibers in order to match their curvatures. By flagging the nodes corresponding to axonal terminals, we were able to accurately calculate axon length, terminal density, and distances between terminals. Rare axonal fibers that mainly innervated dendritic shafts were excluded from analysis because they are presumed to be from interneurons. The exact coordinates were extracted with the VAST API function *getannoobject*, which provides a matrix with all x, y, and z coordinates of every node. We then designed MATLAB scripts that use these matrix coordinates to calculate distances depending on the type of analysis. All distance and length measurements were based on the MATLAB function *vecnorm* that uses the Euclidean norm, where vector v with N elements is defined by:

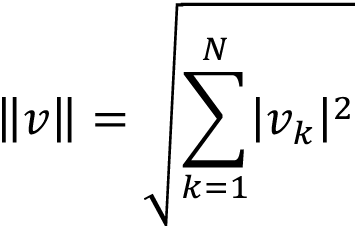

#### 3D Modeling

Final 3D modeling was performed in either the included VAST 3D viewer or in Blender 3.5 (http://Blender.org), an open-source 3D computer-graphics software toolset with modeling, material editing, and rendering capabilities. The VAST 3D viewer was used to visualize the raw EM block and large-saturated reconstructions of axons and dendrites. This was done by combining CDeep3M/PyTorch Connectomics prediction maps with segmentation import functions in VAST. All other modeling was done in Blender. To begin, 3D surface meshes of neuropil structures were imported as .obj files. No post-import size scaling was applied. To reduce computational load, most features were exported at a lower resolution (Mipmap level 2-3). The lower resolution models maintained their native scale. Additionally, importing .obj models from VastTools preserves the spatial location of every segmented feature, so the actual spatial distribution from the SBEM image stack is maintained. Synaptic vesicles were represented by pre-made 40 nm 3D models included with VastTools. Smoothing, color enhancements, and material assignments were applied equally to all conditions and brains. The color-coding and patterning schemes for each feature are included in the figures. All structures except for vesicles were smoothened with the “smooth vertices” function. The scenes were rendered using the cycles renderer.

### Data blinding

To ensure consistency of annotations and to eliminate any bias, manual tracing of all subcellular structures, classifications, and extractions of various synaptic parameters were performed by several trained investigators who were blinded to experimental conditions.

### Quantifications and statistics

All final quantifications, curve fittings, and statistical analyses were performed in Origin Pro. We found that the distributions of most measured parameters, such as terminal/spine distances and sizes of various structures in individual synapses, were significantly deviated from normal. Therefore, non-parametric statistical tests were used throughout most of the study. Standard comparisons of populations (lognormal fitting curve plots) were assessed using Mann-Whitney U Test. For comparison of mice (box with data overlap plots), the *p* values were determined by t-test. Correlation analyses were performed using Spearman or Pearson tests. Statistical significances of differences between Spearman correlation coefficients (rs) were determined with Fisher rs-to-z transformation test. Our choice of *Lognormal probability density* function (PDF) ^67^ for analysis of distributions was based on systematic side-by-side fitting of each dataset with *Gaussian*, *Weibull*, *Lognormal* and *Gamma* fitting functions accompanied with Kolmogorov-Smirnov test for goodness of fits. These analyses indicated that Lognormal PDF was the most versatile.

The PDF of a Lognormal distribution is:

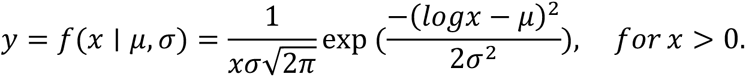

## Supplementary Figures with legends

**Figure S1.**
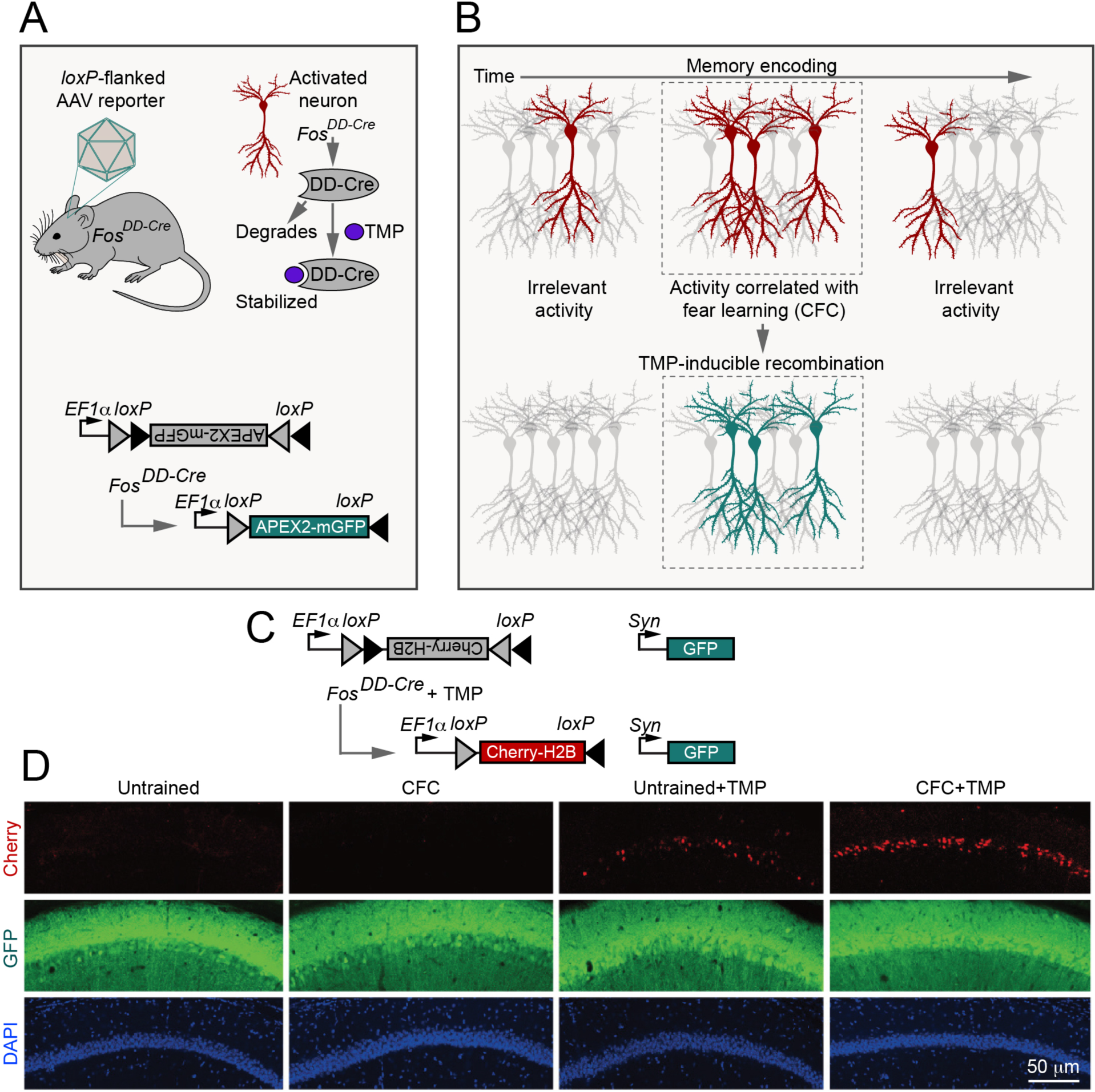
Irreversible chemogenetic labeling of transiently activated neurons with FosDD-Cre. **(A)** Schematics of viral delivery of *loxP*-flanked reporters into the brain of *Fos^DD-Cre^* knock-in mice, TMP-inducible stabilization of DD-Cre in Fos-positive activated neurons, and the DIO:APEX2-mGFP adeno-associated virus (AAV) used for SBEM reconstructions. **(B)** Schematics of acute induction of viral reporters with TMP in neurons recruited for memory acquisition. **(C** and **D)** Sparce experience-dependent labeling of cellular ensembles in *Fos^DD-Cre^* mice does not reflect a poor virus penetrance or recombination efficiency. **(C)** Simultaneous expression of two AAV-driven reporters, Cherry-H2B and GFP, in a Cre-inducible and constitutive manner. **(D)** Confocal images of Cherry, GFP, and DAPI fluorescence in the CA1 of untrained and fear-conditioned mice. Single doses of TMP (50 µg/g body weight) or a control vehicle solution were administered intraperitoneally 30 minutes post-training. Note that Cherry-positive cells are only detected after TMP treatment and their numbers are increased after CFC, whereas the constitutive expression of GFP under the control of Synapsin (Syn) promoter is widespread in all experimental settings. Scale bar applies to all panels.

**Figure S2.**
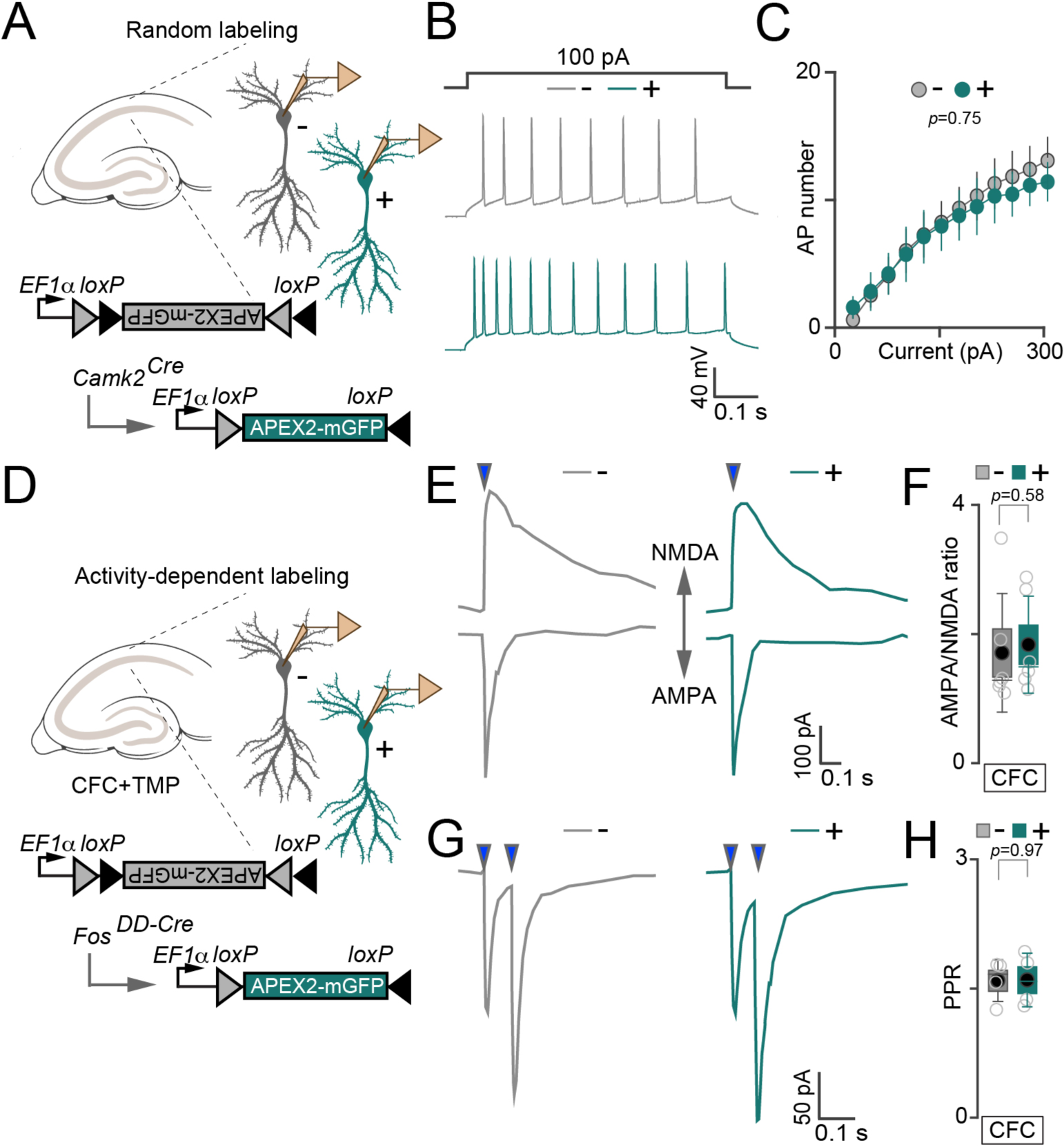
Extended electrophysiological analysis of neuronal excitability and synaptic strength. **(A** to **C)** Control experiments demonstrating that altered electrical properties of PNs with a remote history of activity during associative learning, as summarized in the main Figures 1F to 1H, are not attributed to expression of APEX2-mGFP per se. **(A)** Schematics of side-by-side whole-cell recordings from randomly labeled and neighboring unlabeled CA1 PNs in acute hippocampal slices. Random labeling was achieved by injecting a diluted virus into the hippocampus of constitutive *Camk2^Cre^*driver mice, which mediates recombination in all mature glutamatergic neurons. **(B)** Sample traces of action potentials (APs) evoked by a 100 pA current injection. **(C)** Numbers of APs (Mean ± S.E.M.), plotted relative to stimulus intensity. *n* =3 mice/7 neurons per group. **(D** to **H)** Synaptic properties of CA1 PNs were assessed 7 days after experience-dependent labeling. **(D)** Schematics of side-by-side whole-cell recordings from Fos/APEX2-mGFP-negative (-) and positive (+) cells in acute hippocampal slices from fear-conditioned TMP-treated *Fos^DD-Cre^* mice. **(E** and **F)** PNs with a remote history of activity during associative learning do not exhibit global changes in synaptic strength. Panels show sample traces of evoked AMPA- and NMDA-type excitatory postsynaptic currents (E, blue arrows indicate the times of 1 ms electrical stimulation through a local extracellular electrode) and quantifications of AMPA/NMDA ratios (F). *n* =3 mice/6 neurons per group. In (F) and subsequent similar panels, graphs display individual data points (open circles), mean values (filled circles), standard errors (boxes), standard deviations (vertical whiskers), and medians (horizontal lines). The *p* values were determined by t-test. **(G** and **H)** PNs with a remote history of activity during associative learning do not exhibit global changes in the presynaptic release probability. Panels show sample traces of AMPA-type currents evoked by two closely spaced 1 ms stimuli (G) and quantifications of paired pulse EPSC ratios (PPR). *n* =2 mice/4 neurons per group. In panels (E) and (G), scale bars apply to all traces.

**Figure S3.**
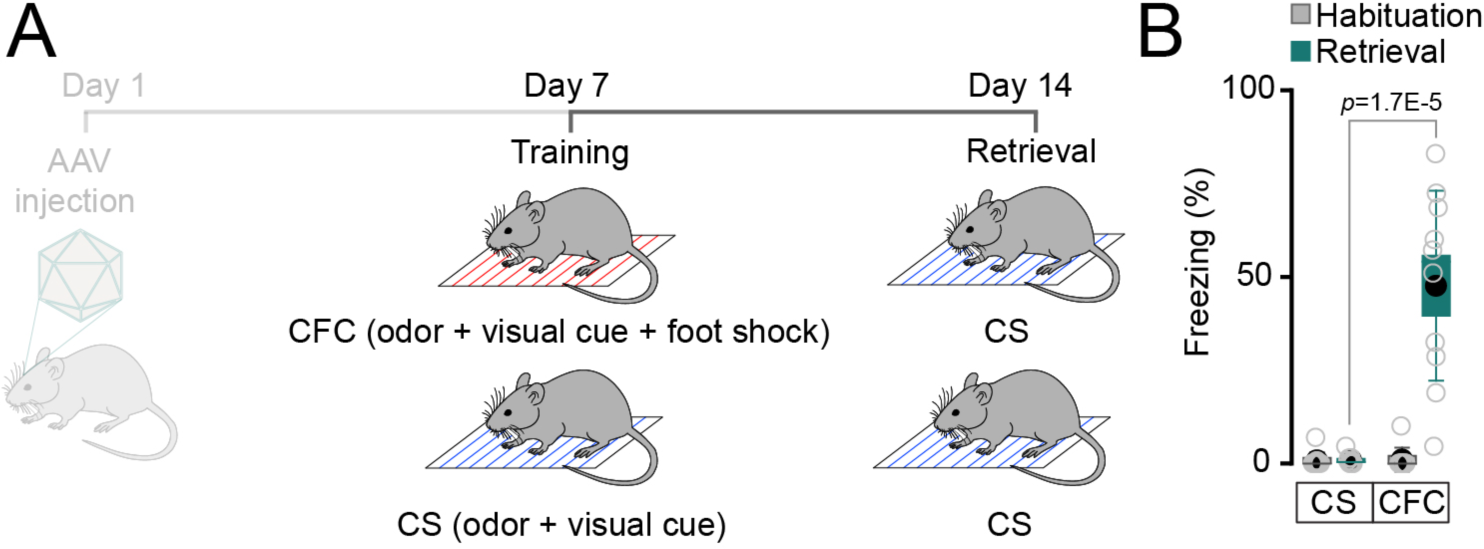
Fear conditioned mice had retrievable memory at the time of tissue collection for SBEM. **(A)** Experimental design. Mice were subjected to CFC or a neutral CS and then re-tested in the original context 7 days later. **(B)** Quantifications of freezing before (habituation) and 7 days after training. *n* = 10 mice per group. The *p* value was determined by t-test.

**Figure S4.**
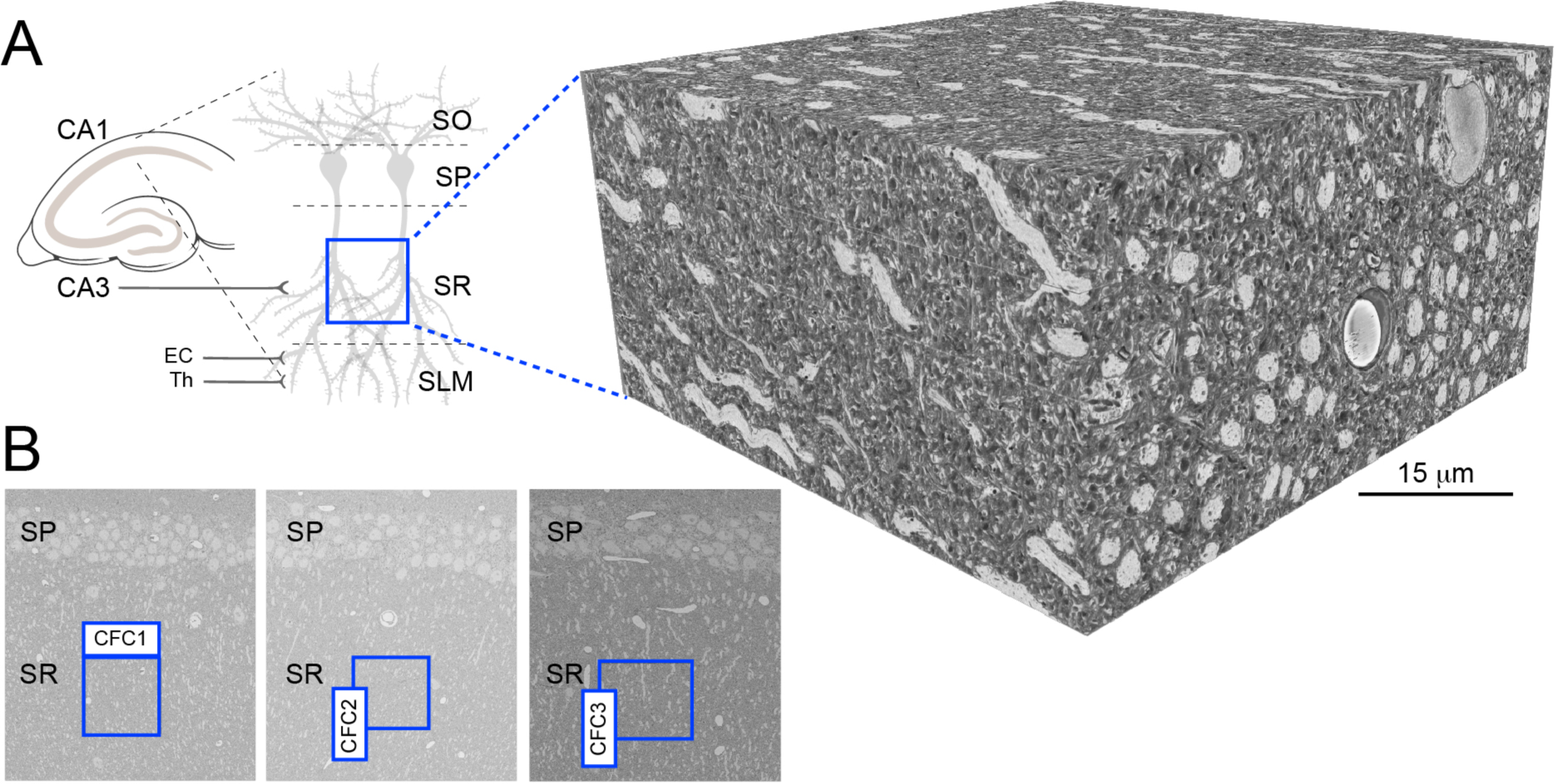
Examples of raw SBEM image data. **(A)** Schematics of excitatory pathways in the CA1 and typical 3D-EM stack collected from the area of stratum radiatum (SR) marked by a blue box. **(B)** Original 2D EM images with actual coordinates of 3D stacks acquired from three different fear conditioned mice (blue boxes).

**Figure S5.**
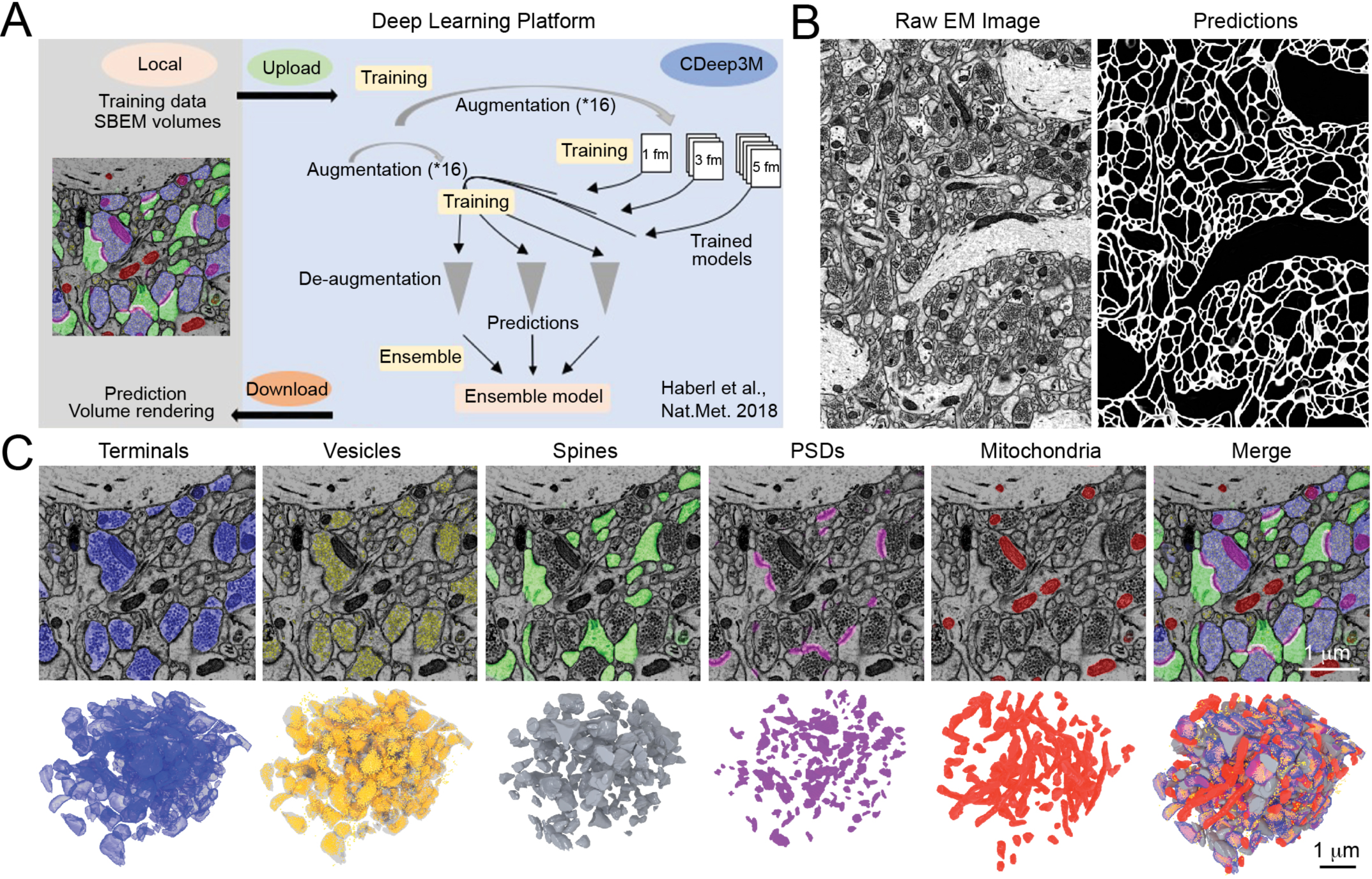
Automated segmentation of subcellular structures in SBEM stacks. **(A)** Workflow for image segmentation in the Amazon cloud-based machine learning platform, CDeep3M. **(B)** Example of automatically segmented plasma membranes. **(C)** Examples of automatically segmented presynaptic terminals, neurotransmitter vesicles, dendritic spines, postsynaptic densities (PSDs), and mitochondria. For each structure, both original 2D EM images with color-coded templates and resulting 3D reconstructions are shown. Scale bars apply to all panels.

**Figure S6.**
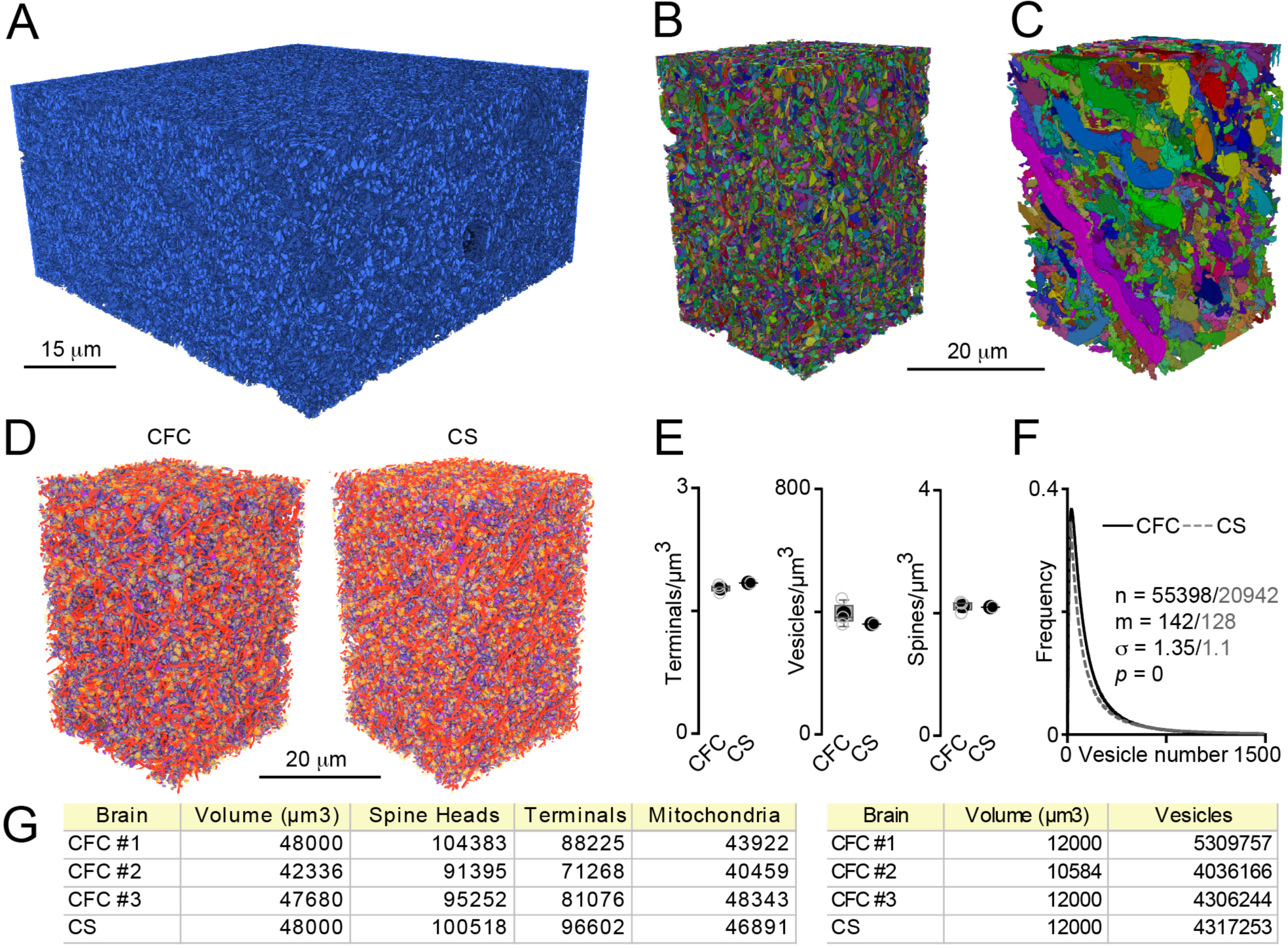
Examples of annotated SBEM stacks with basic quantifications. **(A)** Saturated reconstructions of axons and presynaptic terminals in the entire volume. **(B** and **C)** Saturated reconstructions of axons with presynaptic terminals (B) and dendrites (C) in ∼1/12^th^ of the total 3D volume. Individual projections are displayed in different colors, scale bar applies to both panels. **(D)** Reconstructions of all synapses and mitochondria in volumes acquired from mice subjected to CFC or CS. Structures are color-coded as shown in Figure S5C. Scale bar applies to both images. **(E)** Averaged counts of terminals, vesicles, and spines per μm^3^, as assessed in difference mice. **(F)** Distributions of vesicle counts in individual synapses. Lognormal curves with fitting parameters are shown. **(G)** Raw values for indicated parameters, as assessed in partial volumes.

**Figure S7.**
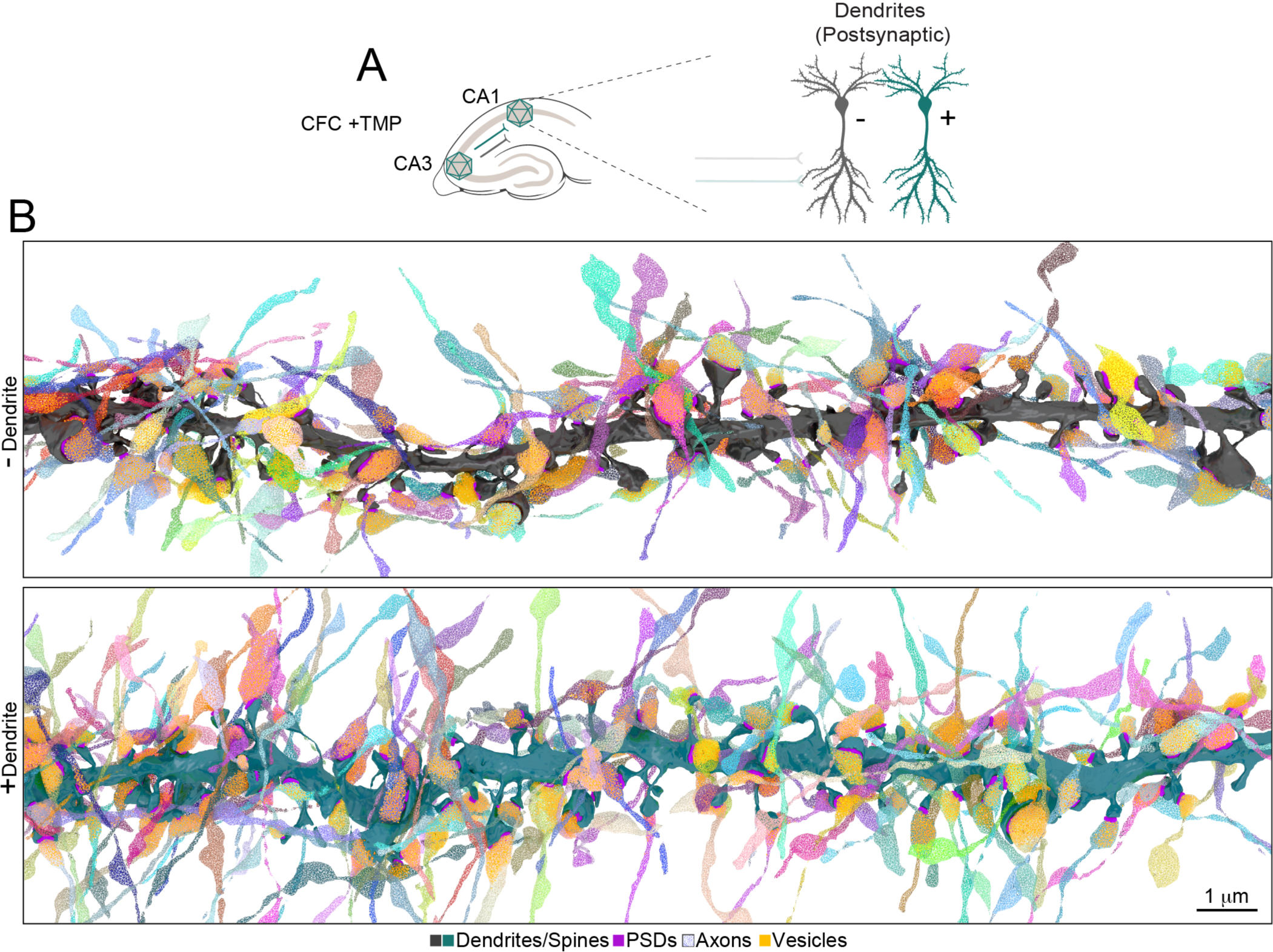
Examples of local connectomes of PNs in the CA1sr of fear conditioned mice. **(A)** Schematics of experience-dependent labeling and data categorization. **(B)** Reconstructions of dendritic branches of Fos/APEX2-mGFP-negative (-) and positive (+) PNs and fibers/terminals of all incoming SchC axons, displayed in different colors.

**Figure S8.**
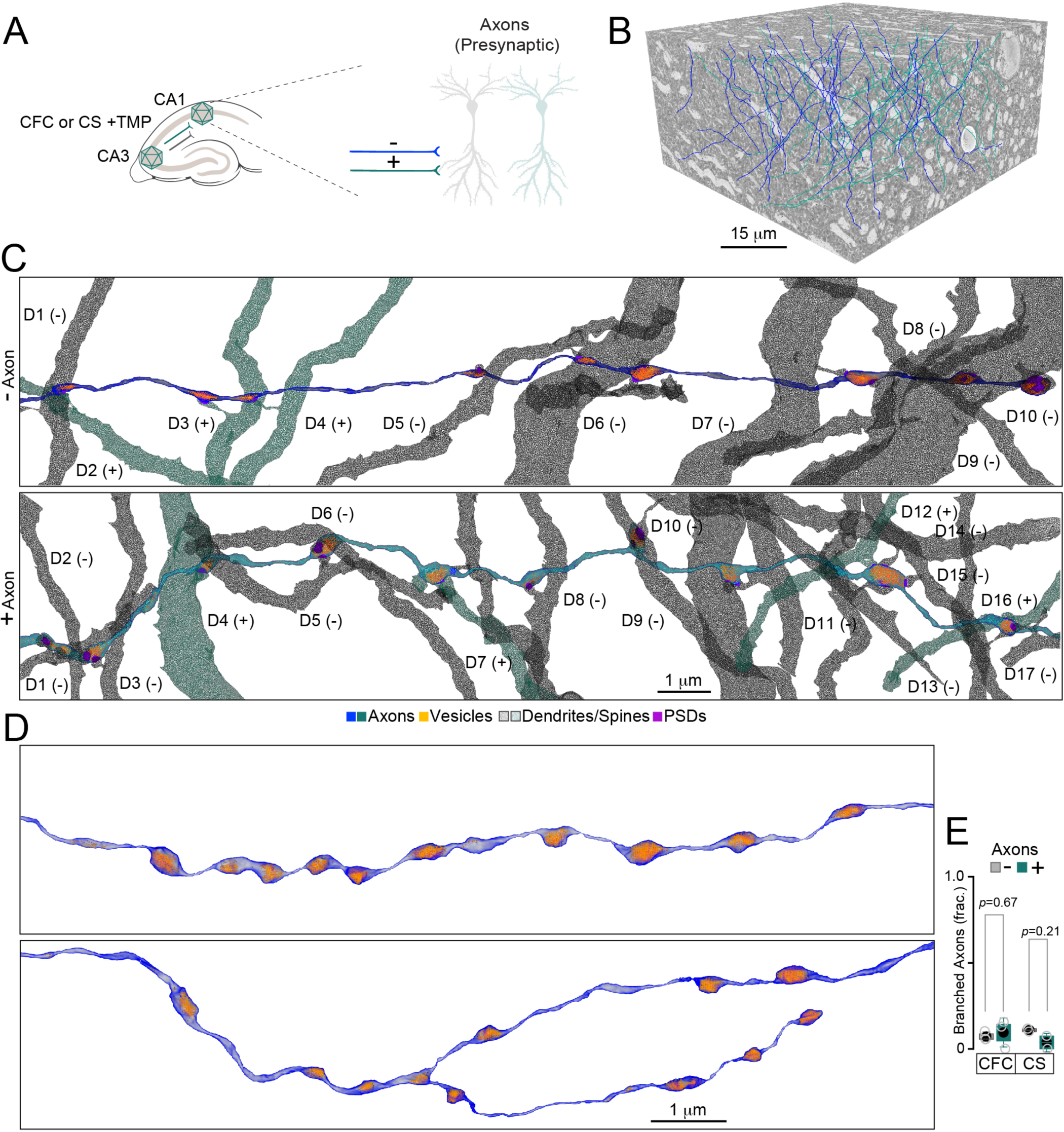
Extended analysis of axonal wiring in the CA1sr of mice subjected to CFC or CS. **(A)** Schematics of experience-dependent labeling and data categorization. **(B)** Typical examples of reconstructed Fos/APEX2-mGFP-negative (-) and positive (+) SchC axons in a full SBEM stack. Only a few unlabeled fibers are displayed. **(C)** Reconstructions of SchC axons terminating in the CA1sr. Different dendrites (marked as D) of CA1 neurons are numbered and color-coded based on their activity history (e.g. Fos/APEX2-mGFP reporter expression). **(D)** Examples of unbranched and branched axons. **(E)** Fractions of branched axons for indicated experimental settings, as assessed in different mice. CFC, *n* = 3 mice; CS, *n* = 2. The *p* values were determined by t-test.

**Figure S9.**
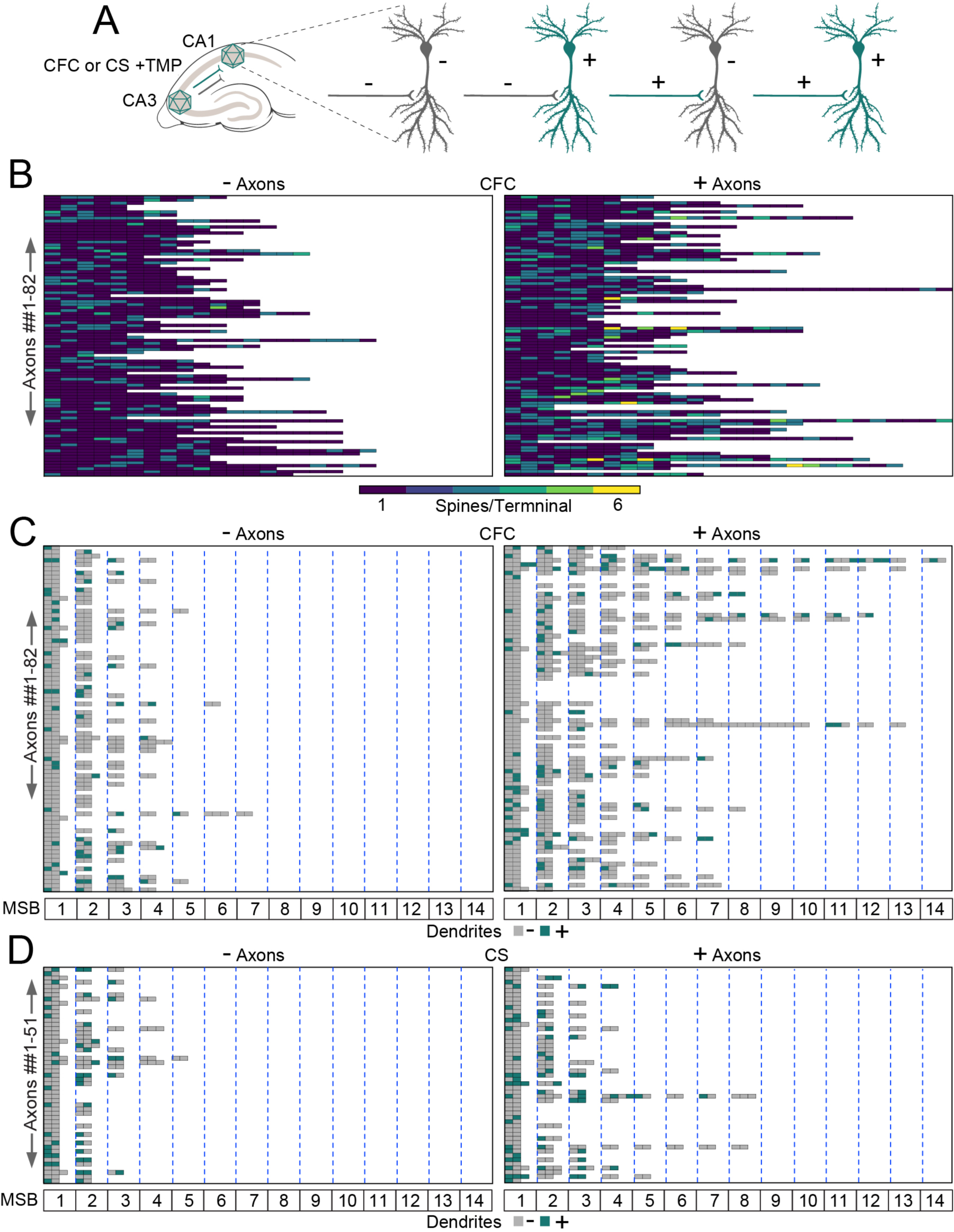
Extended analysis of axonal wiring through MSBs. **(A)** Schematics of experience-dependent labeling and data categorization. **(B)** Heatmaps representing the numbers of spines innervated by each terminal of individual Fos/APEX2-mGFP-negative (-) and positive (+) SchC axon in the CA1sr of fear conditioned mice (*n* = 3). **(C** and **D)** Heatmaps representing axonal wiring via MSBs in the CA1sr of mice subjected to CFC (C, *n* = 3) or CS alone (D, *n* = 2). Each box in the vertical columns shows the numbers of color-coded postsynaptic counterparts of each MSB formed by individual Fos/APEX2-mGFP-negative (-) and positive (+) SchC axons (horizontal rows, SSBs are omitted).

**Figure S10.**
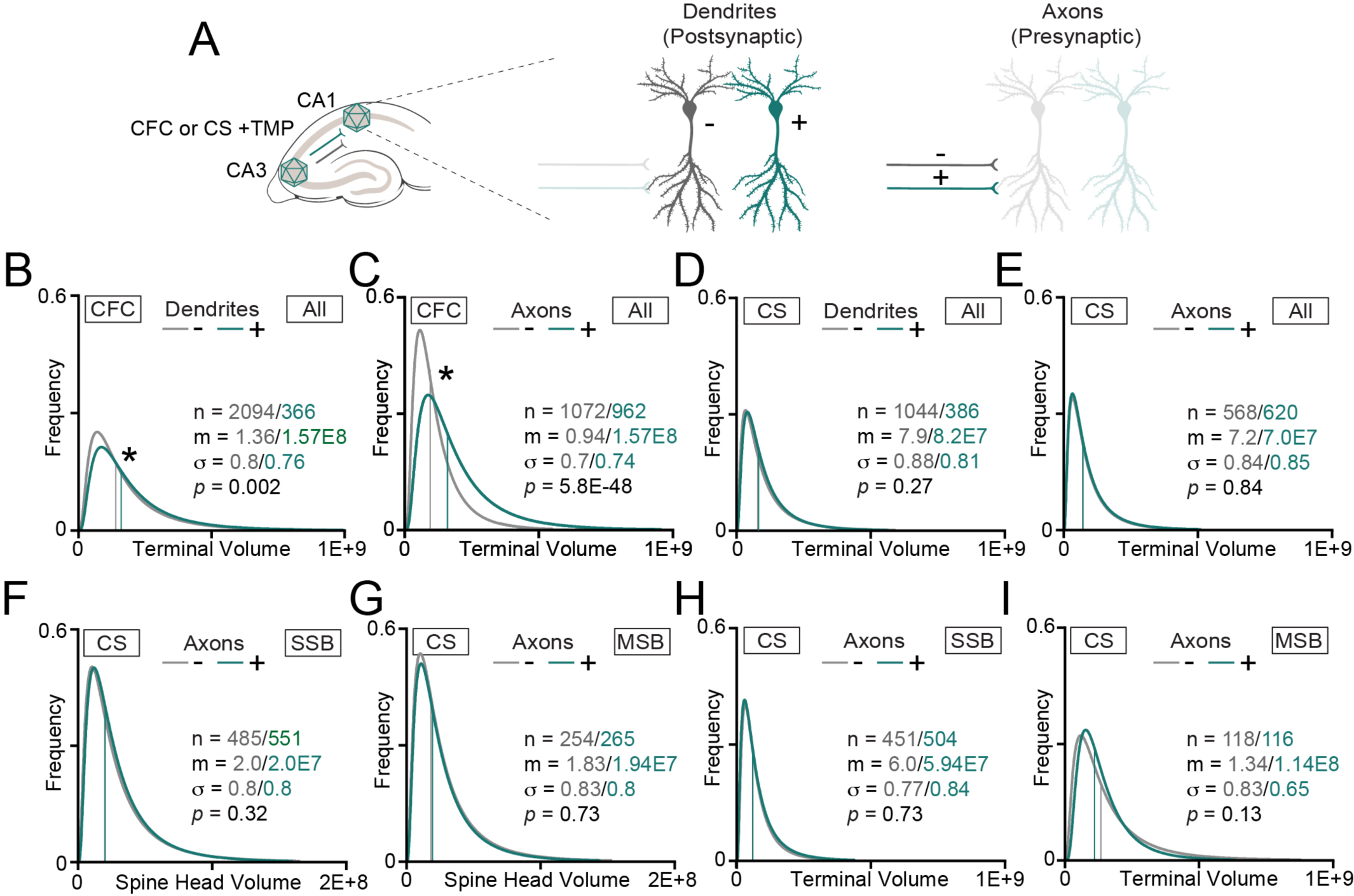
Extended extrapolations of synaptic weights. Sizes of individual excitatory synapses of PNs with remote histories of activity correlated with fear learning (*n* = 3 mice) or exposure to a neutral CS (*n* = 2) were examined in CA1sr using Fos/APEX2-mGFP labeling on either post- (CA1 dendrites) or presynaptic side (SchC axons) as a frame of reference. All volumetric measurements are displayed in nm^3^. **(A)** Schematics of experience-dependent labeling and data categorization. **(B** to **I)** Distributions of terminal and spine head volumes for indicated conditions. Lognormal curves with fitting parameters are shown. n – sample sizes; m – medians; σ - standard deviations of logarithmic values. The *p* values were calculated by Mann-Whitney tests. **(B** and **C)** Volumes of all terminals in fear conditioned mice, categorized by post- (B) or presynaptic labeling (C). **(D** and **E)** Measurements of the same parameters as in (B and C) in mice subjected to a neutral CS. **(F** to **I)** Analysis of conventional SSB and MSB-type synapses in mice subjected to a neutral CS. Datasets were categorized by history of presynaptic activity. **(F)** Head volumes of spines innervated by SSBs. **(G)** Head volumes of spines innervated by MSBs. **(H)** SSB terminal volumes. **(I)** MSB terminal volumes.

**Figure S11.**
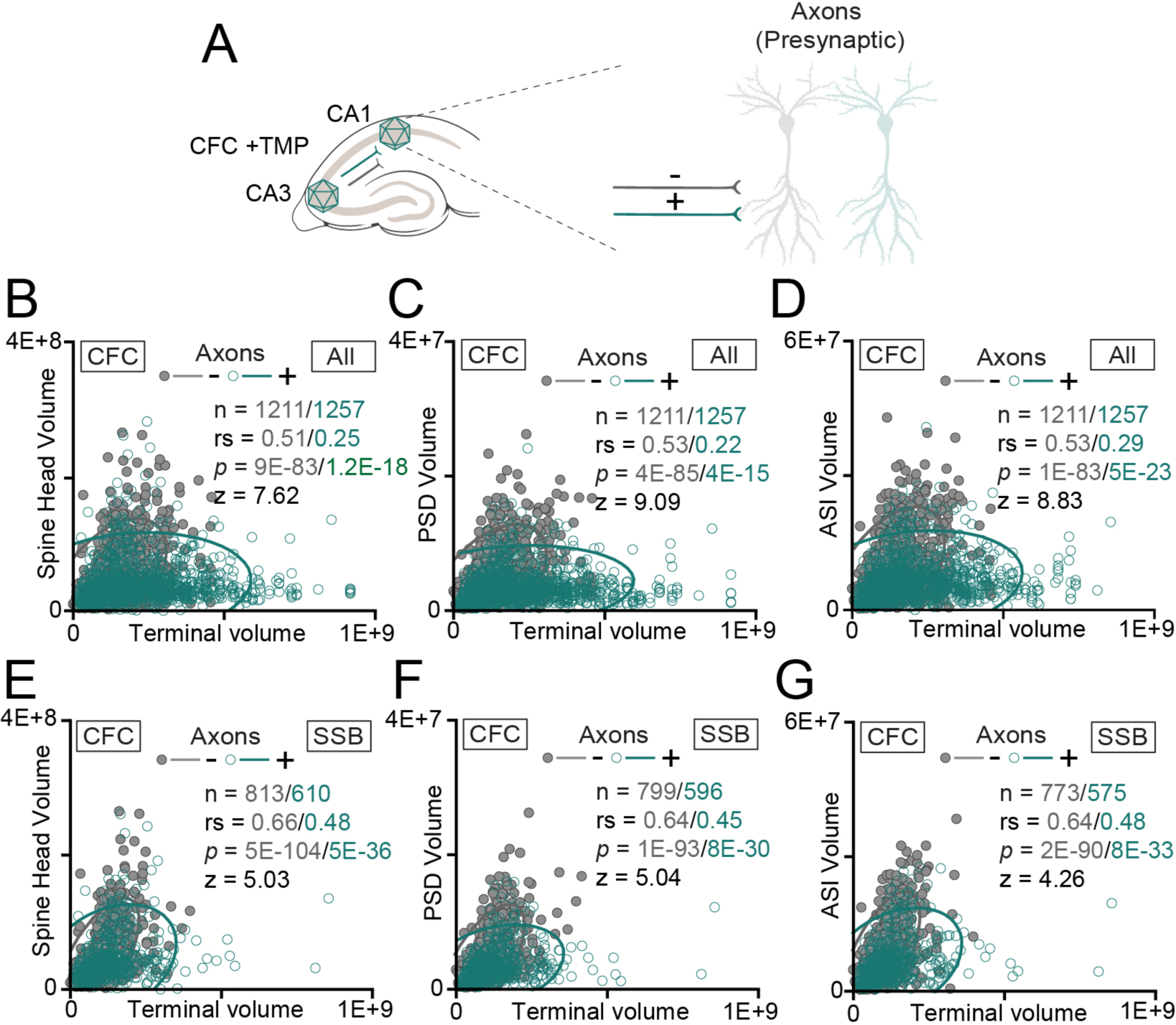
Extended analysis of single synapses. Proportionalities between the sizes of pre- and postsynaptic structures in all or SSB-type excitatory synapses were assessed in mice subjected to CFC (*n* = 3). Datasets were annotated based on the history of CA3 neuron activity (e.g. axonal labeling with Fos/APEX2-mGFP). All volumetric measurements are displayed in nm^3^. **(A)** Schematics of experience-dependent labeling and data categorization. **(B** to **G)** Correlations between indicated parameters. Scatter plots with confidence ellipses, sample sizes (n), Spearman correlation coefficients (rs), Fisher transformation scores (z) and *p* values are shown. **(B)** Terminal vs spine head volumes for all synapses. **(C)** Terminal vs PSD volumes for all synapses. **(D)** Terminal vs ASI volumes for all synapses. **(E)** Terminal vs spine head volumes in SSB-type synapses. **(F)** Terminal vs PSD volumes in SSB-type synapses. **(G)** Terminal vs ASI volumes in SSB-type synapses.

**Figure S12.**
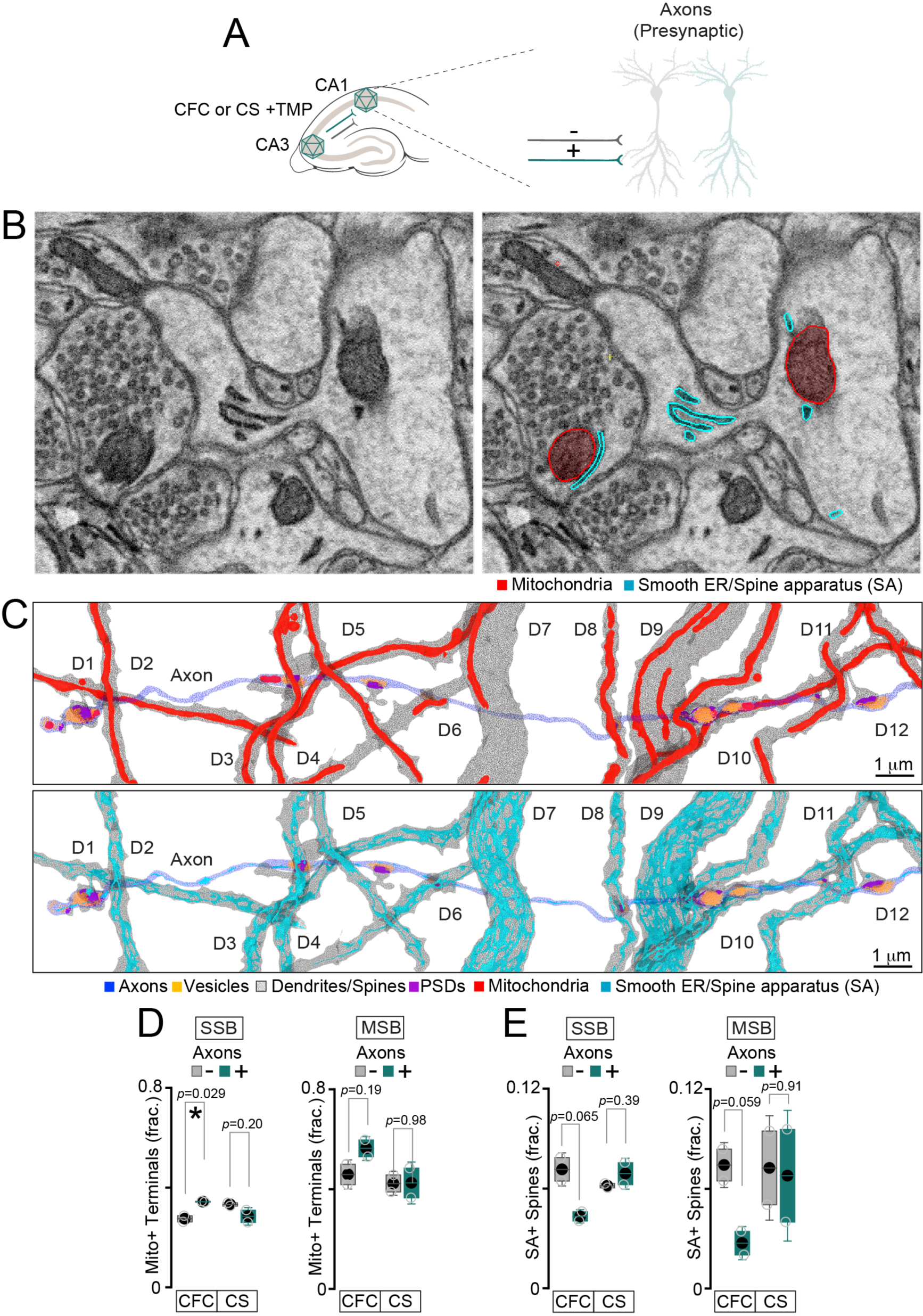
Extended analysis of intracellular membrane organelles. **(A)** Schematics of experience-dependent labeling and data categorization. **(B)** Examples of raw 2D images with traced presynaptic mitochondria (red) and postsynaptic spine apparatus (SA, green). **(C)** Reconstructions of all mitochondria, smooth endoplasmic reticulum (ER) and SA in SchC axons and/or their target dendrites of PNs in CA1sr. Structures are colored as indicated in the legend. **(D)** Fractions of mitochondria-containing SSBs and MSBs formed by Fos/APEX2-mGFP-negative (-) and positive (+) axons of mice subjected to CFC (*n* = 3) or CS alone (*n* = 2). **(E)** Fractions of SA-containing spines in SSB- and MSB-type synapses formed by Fos/APEX2-mGFP-negative (-) and positive (+) axons of mice subjected to CFC (*n* = 3) or CS (*n* = 2). The *p* values were calculated by t-test.

## References

1. Holtmaat, A., and Svoboda, K. (2009). Experience-dependent structural synaptic plasticity in the mammalian brain. Nat Rev Neurosci 10, 647–658. 10.1038/nrn2699.

2. Josselyn, S.A., and Tonegawa, S. (2020). Memory engrams: Recalling the past and imagining the future. Science 367. 10.1126/science.aaw4325.

3. Berry, K.P., and Nedivi, E. (2017). Spine Dynamics: Are They All the Same? Neuron 96, 43–55. 10.1016/j.neuron.2017.08.008.

4. Caroni, P., Donato, F., and Muller, D. (2012). Structural plasticity upon learning: regulation and functions. Nat Rev Neurosci 13, 478–490. 10.1038/nrn3258.

5. Magee, J.C., and Grienberger, C. (2020). Synaptic Plasticity Forms and Functions. Annu Rev Neurosci 43, 95–117. 10.1146/annurev-neuro-090919-022842.

6. Lisman, J. (2017). Glutamatergic synapses are structurally and biochemically complex because of multiple plasticity processes: long-term potentiation, long-term depression, short-term potentiation and scaling. Philos Trans R Soc Lond B Biol Sci 372. 10.1098/rstb.2016.0260.

7. Fox, K., and Stryker, M. (2017). Integrating Hebbian and homeostatic plasticity: introduction. Philos Trans R Soc Lond B Biol Sci 372. 10.1098/rstb.2016.0413.

8. Reijmers, L.G., Perkins, B.L., Matsuo, N., and Mayford, M. (2007). Localization of a stable neural correlate of associative memory. Science 317, 1230–1233. 10.1126/science.1143839.

9. Liu, X., Ramirez, S., Pang, P.T., Puryear, C.B., Govindarajan, A., Deisseroth, K., and Tonegawa, S. (2012). Optogenetic stimulation of a hippocampal engram activates fear memory recall. Nature 484, 381–385. 10.1038/nature11028.

10. Kitamura, T., Ogawa, S.K., Roy, D.S., Okuyama, T., Morrissey, M.D., Smith, L.M., Redondo, R.L., and Tonegawa, S. (2017). Engrams and circuits crucial for systems consolidation of a memory. Science 356, 73–78. 10.1126/science.aam6808.

11. Cowansage, K.K., Shuman, T., Dillingham, B.C., Chang, A., Golshani, P., and Mayford, M. (2014). Direct reactivation of a coherent neocortical memory of context. Neuron 84, 432–441. 10.1016/j.neuron.2014.09.022.

12. Choi, J.H., Sim, S.E., Kim, J.I., Choi, D.I., Oh, J., Ye, S., Lee, J., Kim, T., Ko, H.G., Lim, C.S., and Kaang, B.K. (2018). Interregional synaptic maps among engram cells underlie memory formation. Science 360, 430–435. 10.1126/science.aas9204.

13. Lee, C., Lee, B.H., Jung, H., Lee, C., Sung, Y., Kim, H., Kim, J., Shim, J.Y., Kim, J.I., Choi, D.I., et al. (2023). Hippocampal engram networks for fear memory recruit new synapses and modify pre-existing synapses in vivo. Curr Biol 33, 507–516 e503. 10.1016/j.cub.2022.12.038.

14. Driscoll, L.N., Duncker, L., and Harvey, C.D. (2022). Representational drift: Emerging theories for continual learning and experimental future directions. Curr Opin Neurobiol 76, 102609. 10.1016/j.conb.2022.102609.

15. Rubin, A., Geva, N., Sheintuch, L., and Ziv, Y. (2015). Hippocampal ensemble dynamics timestamp events in long-term memory. Elife 4. 10.7554/eLife.12247.

16. Ziv, Y., Burns, L.D., Cocker, E.D., Hamel, E.O., Ghosh, K.K., Kitch, L.J., El Gamal, A., and Schnitzer, M.J. (2013). Long-term dynamics of CA1 hippocampal place codes. Nat Neurosci 16, 264–266. 10.1038/nn.3329.

17. Driscoll, L.N., Pettit, N.L., Minderer, M., Chettih, S.N., and Harvey, C.D. (2017). Dynamic Reorganization of Neuronal Activity Patterns in Parietal Cortex. Cell 170, 986–999 e916. 10.1016/j.cell.2017.07.021.

18. Schoonover, C.E., Ohashi, S.N., Axel, R., and Fink, A.J.P. (2021). Representational drift in primary olfactory cortex. Nature 594, 541–546. 10.1038/s41586-021-03628-7.

19. Lee, J.S., Briguglio, J.J., Cohen, J.D., Romani, S., and Lee, A.K. (2020). The Statistical Structure of the Hippocampal Code for Space as a Function of Time, Context, and Value. Cell 183, 620–635 e622. 10.1016/j.cell.2020.09.024.

20. Denk, W., and Horstmann, H. (2004). Serial block-face scanning electron microscopy to reconstruct three-dimensional tissue nanostructure. PLoS Biol 2, e329. 10.1371/journal.pbio.0020329.

21. Kasthuri, N., Hayworth, K.J., Berger, D.R., Schalek, R.L., Conchello, J.A., Knowles-Barley, S., Lee, D., Vazquez-Reina, A., Kaynig, V., Jones, T.R., et al. (2015). Saturated Reconstruction of a Volume of Neocortex. Cell 162, 648–661. 10.1016/j.cell.2015.06.054.

22. Morgan, J.L., Berger, D.R., Wetzel, A.W., and Lichtman, J.W. (2016). The Fuzzy Logic of Network Connectivity in Mouse Visual Thalamus. Cell 165, 192–206. 10.1016/j.cell.2016.02.033.

23. Helmstaedter, M., Briggman, K.L., Turaga, S.C., Jain, V., Seung, H.S., and Denk, W. (2013). Connectomic reconstruction of the inner plexiform layer in the mouse retina. Nature 500, 168–174. 10.1038/nature12346.

24. Svara, F.N., Kornfeld, J., Denk, W., and Bollmann, J.H. (2018). Volume EM Reconstruction of Spinal Cord Reveals Wiring Specificity in Speed-Related Motor Circuits. Cell Rep 23, 2942–2954. 10.1016/j.celrep.2018.05.023.

25. Winding, M., Pedigo, B.D., Barnes, C.L., Patsolic, H.G., Park, Y., Kazimiers, T., Fushiki, A., Andrade, I.V., Khandelwal, A., Valdes-Aleman, J., et al. (2023). The connectome of an insect brain. Science 379, eadd9330. 10.1126/science.add9330.

26. Motta, A., Berning, M., Boergens, K.M., Staffler, B., Beining, M., Loomba, S., Hennig, P., Wissler, H., and Helmstaedter, M. (2019). Dense connectomic reconstruction in layer 4 of the somatosensory cortex. Science 366. 10.1126/science.aay3134.

27. Cook, S.J., Jarrell, T.A., Brittin, C.A., Wang, Y., Bloniarz, A.E., Yakovlev, M.A., Nguyen, K.C.Q., Tang, L.T., Bayer, E.A., Duerr, J.S., et al. (2019). Whole-animal connectomes of both Caenorhabditis elegans sexes. Nature 571, 63–71. 10.1038/s41586-019-1352-7.

28. Mishchenko, Y., Hu, T., Spacek, J., Mendenhall, J., Harris, K.M., and Chklovskii, D.B. (2010). Ultrastructural analysis of hippocampal neuropil from the connectomics perspective. Neuron 67, 1009–1020. 10.1016/j.neuron.2010.08.014.

29. Turner, N.L., Macrina, T., Bae, J.A., Yang, R., Wilson, A.M., Schneider-Mizell, C., Lee, K., Lu, R., Wu, J., Bodor, A.L., et al. (2022). Reconstruction of neocortex: Organelles, compartments, cells, circuits, and activity. Cell 185, 1082–1100 e1024. 10.1016/j.cell.2022.01.023.

30. Nakashiba, T., Young, J.Z., McHugh, T.J., Buhl, D.L., and Tonegawa, S. (2008). Transgenic inhibition of synaptic transmission reveals role of CA3 output in hippocampal learning. Science 319, 1260–1264. 10.1126/science.1151120.

31. Dillingham, B., Cameron, P., S, P., Cardozo, L., Yoo, E., Maximov, A., Stowers, L., and Mayford, M. (2019). Fear Learning Induces Long-Lasting Changes in Gene Expression and Pathway Specific Presynaptic Growth. BioRxiv. 10.1101/571331.

32. Sando, R., 3rd, Baumgaertel, K., Pieraut, S., Torabi-Rander, N., Wandless, T.J., Mayford, M., and Maximov, A. (2013). Inducible control of gene expression with destabilized Cre. Nat Methods 10, 1085–1088. 10.1038/nmeth.2640.

33. Yap, E.L., and Greenberg, M.E. (2018). Activity-Regulated Transcription: Bridging the Gap between Neural Activity and Behavior. Neuron 100, 330–348. 10.1016/j.neuron.2018.10.013.

34. Iwamoto, M., Bjorklund, T., Lundberg, C., Kirik, D., and Wandless, T.J. (2010). A general chemical method to regulate protein stability in the mammalian central nervous system. Chem Biol 17, 981–988. 10.1016/j.chembiol.2010.07.009.

35. Guenthner, C.J., Miyamichi, K., Yang, H.H., Heller, H.C., and Luo, L. (2013). Permanent genetic access to transiently active neurons via TRAP: targeted recombination in active populations. Neuron 78, 773–784. 10.1016/j.neuron.2013.03.025.

36. Ye, L., Allen, W.E., Thompson, K.R., Tian, Q., Hsueh, B., Ramakrishnan, C., Wang, A.C., Jennings, J.H., Adhikari, A., Halpern, C.H., et al. (2016). Wiring and Molecular Features of Prefrontal Ensembles Representing Distinct Experiences. Cell 165, 1776–1788. 10.1016/j.cell.2016.05.010.

37. Haberl, M.G., Churas, C., Tindall, L., Boassa, D., Phan, S., Bushong, E.A., Madany, M., Akay, R., Deerinck, T.J., Peltier, S.T., and Ellisman, M.H. (2018). CDeep3M-Plug-and-Play cloud-based deep learning for image segmentation. Nat Methods 15, 677–680. 10.1038/s41592-018-0106-z.

38. Zhu, Y., Uytiepo, M., Bushong, E., Haberl, M., Beutter, E., Scheiwe, F., Zhang, W., Chang, L., Luu, D., Chui, B., et al. (2021). Nanoscale 3D EM reconstructions reveal intrinsic mechanisms of structural diversity of chemical synapses. Cell Rep 35, 108953. 10.1016/j.celrep.2021.108953.

39. Buzsaki, G., and Mizuseki, K. (2014). The log-dynamic brain: how skewed distributions affect network operations. Nat Rev Neurosci 15, 264–278. 10.1038/nrn3687.

40. Bartol, T.M., Bromer, C., Kinney, J., Chirillo, M.A., Bourne, J.N., Harris, K.M., and Sejnowski, T.J. (2015). Nanoconnectomic upper bound on the variability of synaptic plasticity. Elife 4, e10778. 10.7554/eLife.10778.

41. Alvarez, V.A., and Sabatini, B.L. (2007). Anatomical and physiological plasticity of dendritic spines. Annu Rev Neurosci 30, 79–97. 10.1146/annurev.neuro.30.051606.094222.

42. Attardo, A., Fitzgerald, J.E., and Schnitzer, M.J. (2015). Impermanence of dendritic spines in live adult CA1 hippocampus. Nature 523, 592–596. 10.1038/nature14467.

43. Sorra, K.E., and Harris, K.M. (1993). Occurrence and three-dimensional structure of multiple synapses between individual radiatum axons and their target pyramidal cells in hippocampal area CA1. J Neurosci 13, 3736–3748. 10.1523/JNEUROSCI.13-09-03736.1993.

44. Rigby, M., Grillo, F.W., Compans, B., Neves, G., Gallinaro, J., Nashashibi, S., Horton, S., Pereira Machado, P.M., Carbajal, M.A., Vizcay-Barrena, G., et al. (2023). Multi-synaptic boutons are a feature of CA1 hippocampal connections in the stratum oriens. Cell Rep 42, 112397. 10.1016/j.celrep.2023.112397.

45. Bloss, E.B., Cembrowski, M.S., Karsh, B., Colonell, J., Fetter, R.D., and Spruston, N. (2018). Single excitatory axons form clustered synapses onto CA1 pyramidal cell dendrites. Nat Neurosci 21, 353–363. 10.1038/s41593-018-0084-6.

46. Toni, N., Buchs, P.A., Nikonenko, I., Bron, C.R., and Muller, D. (1999). LTP promotes formation of multiple spine synapses between a single axon terminal and a dendrite. Nature 402, 421–425. 10.1038/46574.

47. Geinisman, Y., Berry, R.W., Disterhoft, J.F., Power, J.M., and Van der Zee, E.A. (2001). Associative learning elicits the formation of multiple-synapse boutons. J Neurosci 21, 5568–5573. 10.1523/JNEUROSCI.21-15-05568.2001.

48. Hedrick, N.G., Lu, Z., Bushong, E., Singhi, S., Nguyen, P., Magana, Y., Jilani, S., Lim, B.K., Ellisman, M., and Komiyama, T. (2022). Learning binds new inputs into functional synaptic clusters via spinogenesis. Nat Neurosci 25, 726–737. 10.1038/s41593-022-01086-6.

49. Harris, K.M. (2020). Structural LTP: from synaptogenesis to regulated synapse enlargement and clustering. Curr Opin Neurobiol 63, 189–197. 10.1016/j.conb.2020.04.009.

50. Hirabayashi, Y., Kwon, S.K., Paek, H., Pernice, W.M., Paul, M.A., Lee, J., Erfani, P., Raczkowski, A., Petrey, D.S., Pon, L.A., and Polleux, F. (2017). ER-mitochondria tethering by PDZD8 regulates Ca(2+) dynamics in mammalian neurons. Science 358, 623–630. 10.1126/science.aan6009.

51. Kwon, S.K., Sando, R., 3rd, Lewis, T.L., Hirabayashi, Y., Maximov, A., and Polleux, F. (2016). LKB1 Regulates Mitochondria-Dependent Presynaptic Calcium Clearance and Neurotransmitter Release Properties at Excitatory Synapses along Cortical Axons. PLoS Biol 14, e1002516. 10.1371/journal.pbio.1002516.

52. Devine, M.J., and Kittler, J.T. (2018). Mitochondria at the neuronal presynapse in health and disease. Nat Rev Neurosci 19, 63–80. 10.1038/nrn.2017.170.

53. Bell, M., Bartol, T., Sejnowski, T., and Rangamani, P. (2019). Dendritic spine geometry and spine apparatus organization govern the spatiotemporal dynamics of calcium. J Gen Physiol 151, 1017–1034. 10.1085/jgp.201812261.

54. Konietzny, A., Wegmann, S., and Mikhaylova, M. (2023). The endoplasmic reticulum puts a new spin on synaptic tagging. Trends Neurosci 46, 32–44. 10.1016/j.tins.2022.10.012.

55. Smith, H.L., Bourne, J.N., Cao, G., Chirillo, M.A., Ostroff, L.E., Watson, D.J., and Harris, K.M. (2016). Mitochondrial support of persistent presynaptic vesicle mobilization with age-dependent synaptic growth after LTP. Elife 5. 10.7554/eLife.15275.

56. Maximov, A., and Sudhof, T.C. (2005). Autonomous function of synaptotagmin 1 in triggering synchronous release independent of asynchronous release. Neuron 48, 547–554. 10.1016/j.neuron.2005.09.006.

57. Lewis, T.L., Jr., Kwon, S.K., Lee, A., Shaw, R., and Polleux, F. (2018). MFF-dependent mitochondrial fission regulates presynaptic release and axon branching by limiting axonal mitochondria size. Nat Commun 9, 5008. 10.1038/s41467-018-07416-2.

58. Sudhof, T.C. (2018). Towards an Understanding of Synapse Formation. Neuron 100, 276–293. 10.1016/j.neuron.2018.09.040.

59. Zhu, Y., Huang, M., Bushong, E., Phan, S., Uytiepo, M., Beutter, E., Boemer, D., Tsui, K., Ellisman, M., and Maximov, A. (2019). Class IIa HDACs regulate learning and memory through dynamic experience-dependent repression of transcription. Nat Commun 10, 3469. 10.1038/s41467-019-11409-0.

60. Pieraut, S., Gounko, N., Sando, R., 3rd, Dang, W., Rebboah, E., Panda, S., Madisen, L., Zeng, H., and Maximov, A. (2014). Experience-dependent remodeling of basket cell networks in the dentate gyrus. Neuron 84, 107-122. 10.1016/j.neuron.2014.09.012.

61. Sando, R., Bushong, E., Zhu, Y., Huang, M., Considine, C., Phan, S., Ju, S., Uytiepo, M., Ellisman, M., and Maximov, A. (2017). Assembly of Excitatory Synapses in the Absence of Glutamatergic Neurotransmission. Neuron 94, 312–321 e313. 10.1016/j.neuron.2017.03.047.

62. Kremer, J.R., Mastronarde, D.N., and McIntosh, J.R. (1996). Computer visualization of three-dimensional image data using IMOD. J Struct Biol 116, 71–76. 10.1006/jsbi.1996.0013.

63. Schindelin, J., Arganda-Carreras, I., Frise, E., Kaynig, V., Longair, M., Pietzsch, T., Preibisch, S., Rueden, C., Saalfeld, S., Schmid, B., et al. (2012). Fiji: an open-source platform for biological-image analysis. Nat Methods 9, 676–682. 10.1038/nmeth.2019.

64. Lin, Z.W., Donglai; Lichtman, Jeff; Pfister, Hanspeter (2021). PyTorch Connectomics: A Scalable and Flexible Segmentation Framework for EM Connectomics. arXiv.

65. Berger, D.R., Seung, H.S., and Lichtman, J.W. (2018). VAST (Volume Annotation and Segmentation Tool): Efficient Manual and Semi-Automatic Labeling of Large 3D Image Stacks. Front Neural Circuits 12, 88. 10.3389/fncir.2018.00088.

66. Risher, W.C., Ustunkaya, T., Singh Alvarado, J., and Eroglu, C. (2014). Rapid Golgi analysis method for efficient and unbiased classification of dendritic spines. PLoS One 9, e107591. 10.1371/journal.pone.0107591.

67. Mood, A.M., Graybill, F.A., and Boes, D.C. (1973). Introduction to the theory of statistics, 3d Edition (McGraw-Hill).

